# Cell-type-resolved Metabolic Flux Inference Reveals Stromal Metabolic Reprogramming Across Human Cardiomyopathies

**DOI:** 10.64898/2026.04.19.719409

**Authors:** Tomoya Sakuma, Satoshi Ohno, Hideyuki Shimizu

**Author notes:** Corresponding authors: Satoshi Ohno, Hideyuki Shimizu.

## Abstract

Metabolic remodeling is a hallmark of cardiomyopathy, yet which cell types bear the metabolic burden and how cell-type-specific contributions are disrupted remain unclear. Here, we developed a cell-type-resolved genome-scale metabolic flux inference pipeline optimized for post-mitotic cardiac tissue by maximizing ATP synthesis rather than biomass production and applied it to a single-nucleus transcriptomic atlas of human cardiomyopathies (78 donors, 869,449 nuclei). Metabolic impairment in dilated cardiomyopathy (DCM) was most profound in stromal cells, whereas myeloid cells exhibited opposing metabolic activation. DCM-associated impairment followed a genotype-dependent severity gradient from structural gene mutations to pathogenic variant-negative (PVneg) cases. PVneg hearts uniquely harbored 24 altered metabolic pathways not significant in any other genotype. These PVneg-specific signatures were independent of clinical severity, indicating a genotype-intrinsic metabolic program. Extending the analysis to arrhythmogenic cardiomyopathy and hypertrophic cardiomyopathy showed that ATP depletion is shared across cardiomyopathy subtypes, whereas metabolic remodeling differed across disease subtypes. Additionally, gene regulatory network analysis linked these alterations to broad transcription factor (TF) dysregulation and pervasive TF–metabolic coupling across all cell types. These findings redefine PVneg DCM as a metabolically distinct entity and reveal conserved stromal metabolic remodeling across cardiomyopathies, providing a framework for genotype-informed mechanistic stratification.

## Introduction

Cardiomyopathies constitute a heterogeneous group of myocardial disorders including dilated (DCM)^1^, hypertrophic (HCM)^2^, and arrhythmogenic (ACM)^3^ subtypes, and represent a major cause of heart failure, arrhythmia, and heart transplantation. Among these, DCM is the most prevalent form and a leading indication for heart transplantation. Because the adult heart is among the most energy-demanding organs in the body, it is particularly vulnerable to metabolic perturbations^4^. Accordingly, metabolic remodeling is increasingly recognized not merely as a consequence of pump failure but as a central component of disease pathogenesis^5,6^. Yet, how metabolic remodeling differs at the cellular level across DCM subtypes, including genetically defined forms such as titin (*TTN*)^7^ and lamin A/C (*LMNA*)^8^ mutations, as well as pathogenic variant-negative (PVneg) DCM, which accounts for the majority of cases, remains an open question. Defining how metabolic dysfunction is distributed across cardiac cell types is therefore essential for mechanistic stratification and precision medicine in this population.

Single-nucleus RNA-sequencing (snRNA-seq) studies have transformed our view of the failing heart by revealing coordinated remodeling across cardiomyocytes, fibroblasts, endothelial cells, mural cells, and immune populations, including genotype-dependent shifts in cell composition and the emergence of activated stromal states^9–11^. In parallel, tissue-level metabolic studies have documented energy starvation^12^, depletion of fatty acid intermediates^13^, and a substrate shift toward ketone body utilization in failing hearts^14^. However, these bulk-tissue approaches cannot resolve cell-type-specific metabolic contributions. This is because different cardiac cell types have fundamentally distinct metabolic demands — cardiomyocytes depend on fatty acid oxidation for contractile energy^15^, fibroblasts sustain extracellular matrix turnover^16^, and myeloid cells undergo metabolic reprogramming upon activation^17^. Consequently, which cell types bear the metabolic burden of cardiomyopathy, whether distinct subtypes converge on shared or divergent metabolic programs, and whether genotype-specific metabolic differences exist at cell-type resolution remain unknown. In particular, the metabolic basis of PVneg DCM relative to genetically defined DCM has not been established.

Inferring metabolic flux from transcriptomic data offers a route to bridge this gap because flux inference incorporates metabolic network structure and stoichiometric constraints that cannot be captured by gene expression alone^18^. Genome-scale metabolic models typically calculate fluxes by maximizing a biological objective function, subject to constraints derived from gene expression levels^19^. However, available genome-scale metabolic modeling methods using single-cell gene expressions, including METAFlux^20^ and scFBA^21^, adopt biomass maximization as their objective function, an assumption designed for proliferating cells (e.g. cancer cells) and poorly matched to the post-mitotic adult myocardium, where maintaining ATP homeostasis rather than generating biomass is the dominant metabolic demand^22^. The absence of a cardiac-appropriate objective function has limited rigorous cell-type-resolved flux analysis in cardiomyopathy.

Here, we developed a cell-type-resolved metabolic flux inference pipeline for cardiac tissue that maximizes ATP synthesis rather than biomass production and applied it to a large-scale snRNA-seq atlas of human cardiomyopathies (78 donors, 869,449 nuclei; **Fig. 1**), with validation in two independent cohorts (**Table S1**). This approach revealed pan-cellular ATP depletion, stromal-predominant metabolic

**Figure 1.**
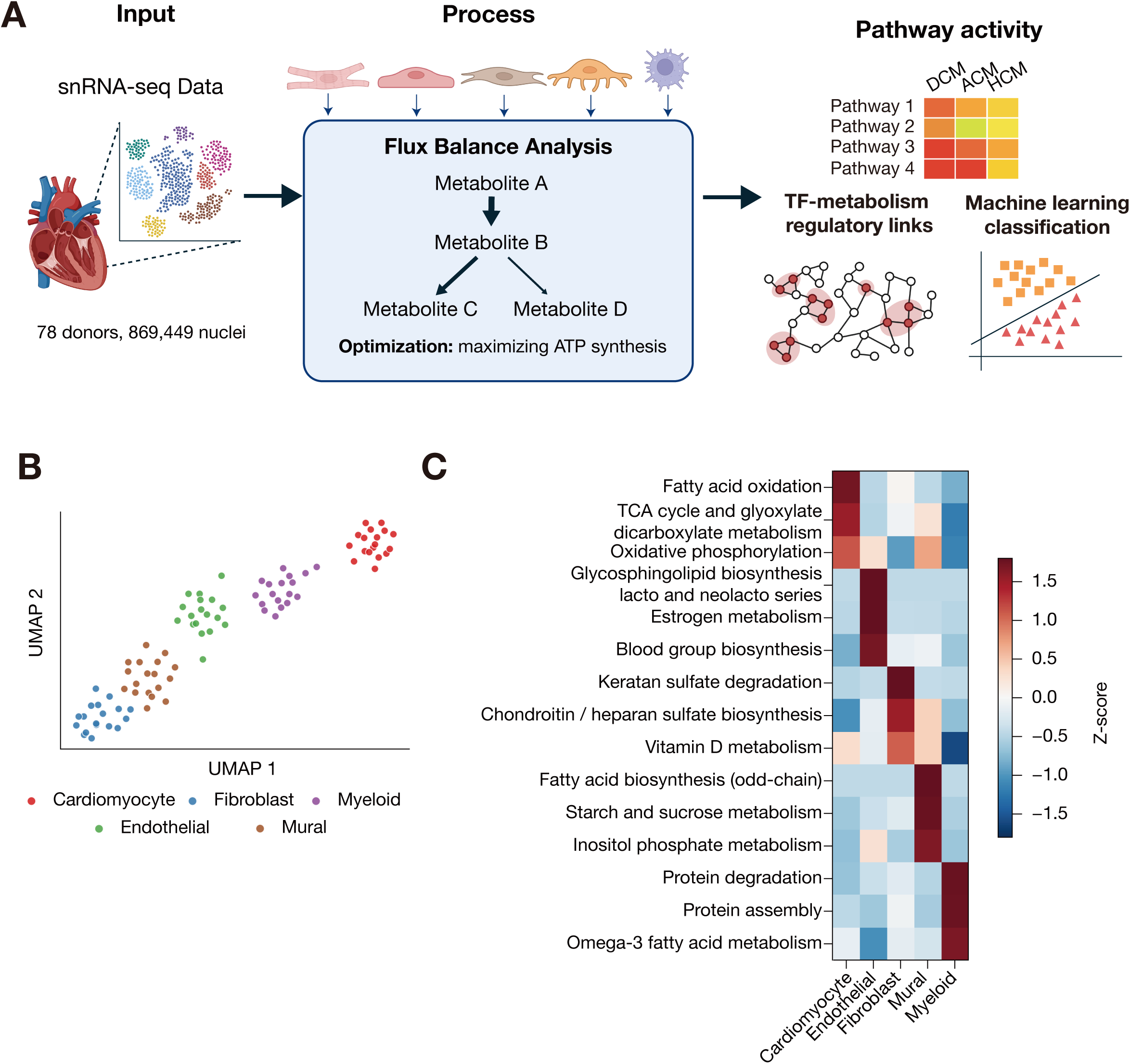
Cell-type-resolved metabolic flux inference and physiological validation. **(A)** Schematic overview of the analysis framework. Single-nucleus RNA-seq (snRNA- seq) profiles from the Human Heart Cell Atlas (n=78 donors) are processed through a cardiac-specific flux balance analysis pipeline that maximizes ATP synthesis under steady-state mass-balance constraints, yielding flux distributions aggregated into 139 metabolic pathway scores. **(B)** UMAP visualization of metabolic pathway profiles in the healthy heart. Pathway scores (139 pathways in Human1^24^) from 18 healthy donors were subjected to principal component analysis followed by UMAP^25^ dimensionality reduction. Each point represents one donor–cell type combination, colored by cell type. **(C)** Physiological validation heatmap. Z-scores for 15 representative pathways across five cell types in healthy donors (n=18). For each pathway, mean absolute flux was computed per cell type and Z-score standardized across the five cell types. Pathways were selected from several biological categories based on published literature. All 15 pathways exhibited the highest Z-score in the expected cell type, confirming physiological validity.

suppression, and genotype-dependent remodeling, and identified PVneg DCM as a metabolically distinct entity characterized by genotype-intrinsic metabolic programs. Cross-disease analyses further indicated that ATP depletion is shared across cardiomyopathy subtypes, while pathway similarity varies by cell type, with fibroblasts showing comparatively conserved profiles across disease comparisons. Additionally, gene regulatory network^23^ analysis further revealed pan-cellular transcription factor dysregulation functionally coupled to metabolic remodeling, despite largely genotype-independent regulon activity. Collectively, these findings provide new insights into the cell-type-specific metabolic organization of cardiomyopathy and a basis for genotype-informed patient stratification.

## Results

### Development and physiological validation of a cell-type-resolved cardiac metabolic flux inference pipeline

The adult heart is a post-mitotic, energy-intensive organ that derives 50–70% of its ATP from fatty acid beta-oxidation, making ATP homeostasis rather than biomass accumulation the dominant metabolic objective of cardiac cells^22^. Existing genome-scale flux inference tools using single-cell gene expressions typically adopt biomass maximization as their objective function, an assumption suited for proliferating cells but misaligned with cardiac physiology^20,21^. We therefore developed a cell-type-resolved metabolic flux inference pipeline optimized for post-mitotic cardiac tissue by maximizing ATP synthesis rather than biomass production (**Fig. 1A**; see Methods) and applied it to 869,449 single-nucleus RNA-sequencing (snRNA-seq) profiles from 78 donors (Reichart cohort^9^; **Table S1**).

To assess whether this pipeline captures biologically meaningful cell-type-specific metabolic identities, activity scores for 139 metabolic pathways defined by Human1^24^, a genome-scale metabolic model (GEM), were computed in healthy donors (n=18) and visualized by UMAP^25^ (**Fig. 1B**). The five major cardiac cell types formed well-separated clusters, with cardiomyocytes and fibroblasts at opposite metabolic poles, indicating that metabolic pathway usage is a robust discriminator of cardiac cell identity.

Flux profiles were then examined against established cell-type biology across 15 pathways selected from cell-type-specific metabolic functions (see Methods; **Fig. 1C**). Each pathway was predicted to be most active in its biologically expected cell

type. Cardiomyocytes exhibited the highest Z-scores for fatty acid oxidation (Z:

+1.70), the TCA cycle (+1.51), and oxidative phosphorylation (+1.11), consistent with their exceptional mitochondrial density and reliance on oxidative metabolism^15,26^. Endothelial cells were enriched for glycosphingolipid biosynthesis of the lacto and neolacto series (+1.79), estrogen metabolism (+1.79), and blood group biosynthesis (+1.70), reflecting glycocalyx maintenance^27^, eNOS-mediated vasodilation^28^, and ABO blood-group antigens^29,30^, respectively. Fibroblasts exhibited high activity in keratan sulfate degradation (+1.79), chondroitin/heparan sulfate biosynthesis (+1.50), and vitamin D metabolism (+1.06), consistent with ECM turnover^16^ and VDR-mediated fibrosis regulation^31^. Mural cells were characterized by inositol phosphate metabolism (+1.65), consistent with IP_3_-mediated Ca^2^⁺ signaling in vascular smooth muscle^32^, and myeloid cells by protein degradation (+1.76), consistent with macrophage antigen processing^33^. The concordance between predicted fluxes and known cell-type physiology supports the biological validity of the inferred metabolic landscape.

To confirm that these physiologically valid profiles were specific to the ATP- centric objective, we benchmarked against conventional biomass maximization^20^. Cardiomyocyte pathway log_2_ fold changes (FC, DCM versus healthy donors) showed low correlation between the two objectives (Spearman ρ = 0.165, *p* = 0.053; **Fig. S1A**), indicating that the two objectives capture fundamentally distinct metabolic landscapes. ATP maximization assigned 28.4% of total ATP synthesis flux to cardiomyocytes, a proportion consistent with their dominant bioenergetic role, whereas biomass maximization distributed flux nearly equally across all cell types (17.9–22.8%; **Fig. S1B**). Applying the 15-pathway validation to biomass-maximized fluxes confirmed this divergence. Biomass maximization achieved only 11/15 correct assignments, misassigning cardiomyocyte energy pathways (fatty acid oxidation, TCA cycle, oxidative phosphorylation) and endothelial blood group biosynthesis (**Fig. S1C**). These results demonstrate that ATP synthesis maximization is the physiologically appropriate objective function for cell-type-resolved metabolic modeling of the adult heart, and that the resulting flux profiles are sufficiently accurate to serve as a foundation for disease analysis.

### ATP depletion across all cell types and asymmetric metabolic remodeling in DCM

Prior bulk-tissue studies have documented reduced ATP production in the failing heart^12^, but whether this energetic deficit is restricted to cardiomyocytes or extends across the cardiac cellular ecosystem has remained unknown. To address this, we first evaluated energetic capacity by computing ATP synthesis flux across all five major cell types in DCM versus healthy donors (**Fig. 2A**). ATP synthesis flux was significantly reduced in all five cell types (*p* < 0.05 for all). Unexpectedly, the most pronounced reductions were observed in stromal cells, with fibroblasts (FC = 0.55, *p* = 0.010), mural cells (FC = 0.58, *p* = 0.005), and endothelial cells (FC = 0.61, *p* = 0.012) showing greater ATP depletion than cardiomyocytes (FC = 0.75, *p* = 0.020), with myeloid cells at intermediate levels. These findings reframe DCM as a pan-cellular energy crisis in which the metabolic burden falls disproportionately on the stromal compartment.

**Figure 2.**
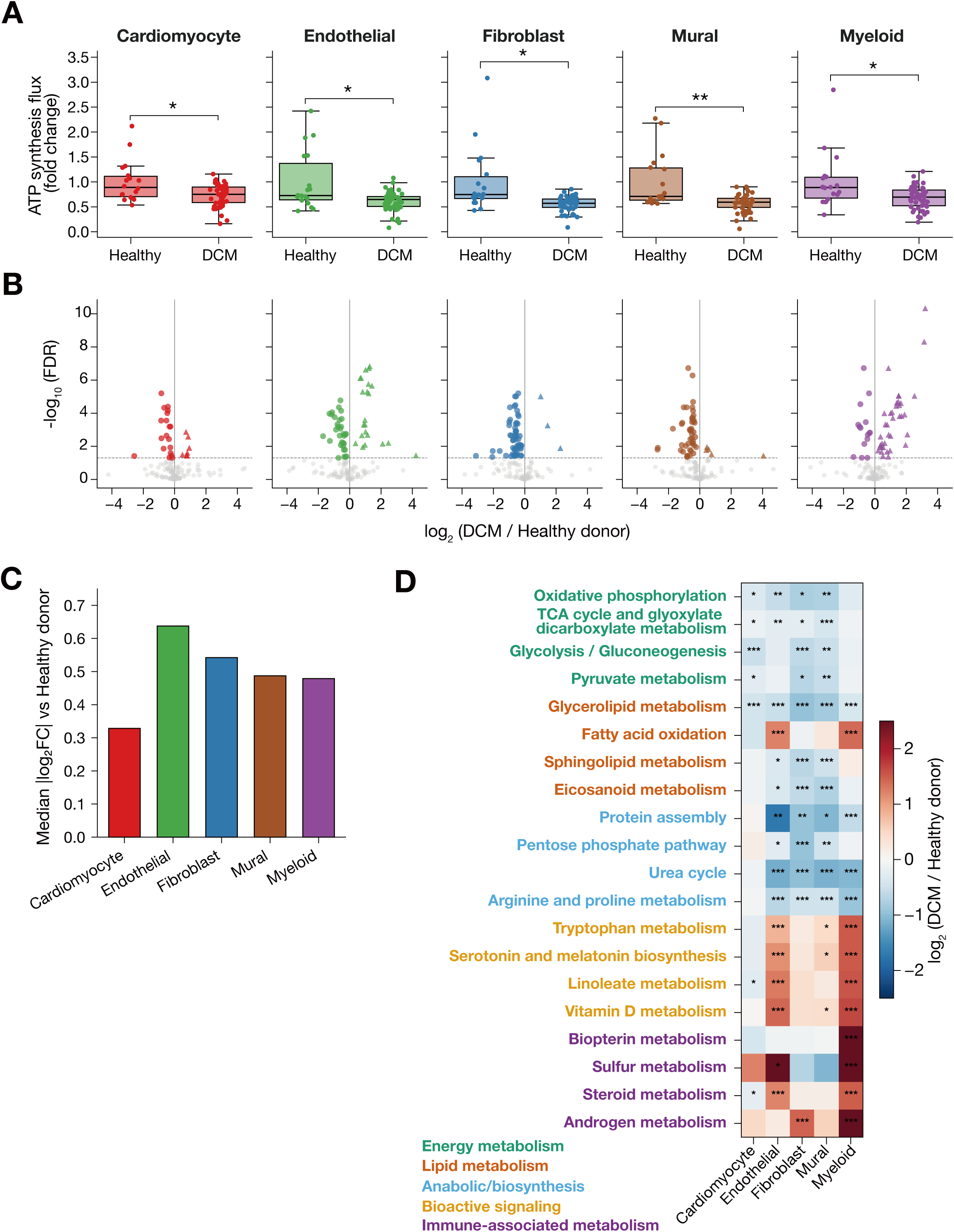
ATP depletion across all cell types and asymmetric metabolic remodeling in DCM. **(A)** ATP synthesis flux fold change across five major cell types in healthy donors (n=18) and DCM (n=52) donors. Values are expressed as fold change relative to the healthy donor mean. Each dot represents one donor. Box plots show median and interquartile range. All five cell types exhibited significant reductions in DCM. Welch’s *t*-test with Benjamini–Hochberg correction: \**p* < 0.05, \*\**p* < 0.01. **(B)** Faceted volcano plots of pathway-level changes (DCM versus healthy donors) for each cell type. Each point represents one of 139 metabolic pathways defined in Human1^24^. Filled triangles indicate pathways significantly upregulated in DCM; filled circles indicate pathways significantly downregulated; gray points are non-significant. Dashed line indicates FDR = 0.05. Fibroblasts and mural cells were dominated by downregulated pathways, whereas myeloid cells exhibited predominantly upregulated pathways. **(C)** Median |log_2_FC| versus healthy donors across 139 metabolic pathways per cell type, computed from DCM donors (n=52) compared with healthy donors (n=18). Cardiomyocytes showed the smallest median effect size, whereas stromal cell types (endothelial, fibroblast, mural) exhibited larger pathway-level changes. **(D)** Curated heatmap of 20 representative metabolic pathways across five cell types, organized by biological category (energy metabolism, lipid metabolism, anabolic/biosynthesis, bioactive signaling, and immune-associated metabolism). Color scale represents log_2_ fold change (DCM versus healthy donors). Asterisks denote statistical significance (\**p* < 0.05, \*\**p* < 0.01, \*\*\**p* < 0.001; FDR-corrected).

This unexpected pattern of preferential stromal ATP depletion prompted us to ask whether the underlying pathway-level remodeling was similarly asymmetric across cell types. Comprehensive analysis of all cell type–pathway combinations identified 247 significantly altered combinations in DCM (FDR < 0.05; **Fig. 2B**), revealing striking directional asymmetry. Fibroblasts showed 59 significantly altered pathways, of which 95% (56/59) were downregulated, and mural cells displayed a similar pattern with 86% (45/52) of altered pathways suppressed, together defining a program of broad metabolic silencing in the stromal compartment. In contrast, myeloid cells exhibited an opposing pattern in which 71% (37/52) of altered pathways were upregulated, indicating immune metabolic activation concurrent with stromal suppression. Consistent with these observations, the magnitude of pathway-level changes (median |log2FC|) was also smaller in cardiomyocytes (0.33) than in stromal cell types (endothelial 0.64, fibroblast 0.54, mural 0.49; **Fig. 2C**), further supporting the conclusion that stromal cells, rather than cardiomyocytes, drive the bulk of pathway-level metabolic remodeling in DCM.

Examination of 20 representative pathways spanning energy metabolism, lipid metabolism, anabolic/biosynthesis, bioactive signaling, and immune-associated metabolism revealed three organizing principles of this asymmetric remodeling (**Fig. 2D**; **Table S2**). First, core energy-producing pathways were broadly suppressed across all five cell types. Oxidative phosphorylation was consistently reduced in cardiomyocytes (log_2_FC: −0.34), endothelial cells (−0.57), fibroblasts (−0.77), and mural cells (−0.74; all FDR < 0.05), providing a direct mechanistic basis for the observed tissue-level ATP depletion. Glycolysis was similarly suppressed in cardiomyocytes (−0.53), fibroblasts (−0.62), and mural cells (−0.51). Second, glycerolipid metabolism was significantly reduced across all five cell types (log2FC: −0.37 to −0.95; all FDR < 0.001), pointing to a pan-cellular lipid metabolic defect that may compromise membrane integrity and lipid signaling in DCM. Third, bioactive signaling pathways — tryptophan metabolism, serotonin and melatonin biosynthesis, linoleate metabolism, and vitamin D metabolism — were markedly upregulated in endothelial and myeloid cells (tryptophan metabolism log_2_FC: +0.83 in endothelial cells, +1.52 in myeloid cells) while trending downward in cardiomyocytes. This directional divergence extended to myeloid-associated pathways such as biopterin, sulfur, steroid, and androgen metabolism, all of which were strongly upregulated in myeloid cells (all FDR < 0.001), reflecting active stress and inflammatory responses within the vascular and immune compartments of the failing heart. Collectively, these results demonstrate that metabolic failure in DCM comprises two opposing programs, with broad energy suppression dominating the stromal compartment and stress-associated metabolic activation defining the immune and vascular lineages.

### Genotype-dependent severity gradient and pathogenic variant-negative-specific metabolic signatures

DCM encompasses a spectrum of genetic etiologies, yet whether distinct causal mutations produce qualitatively different or merely quantitatively varying metabolic phenotypes has not been established at cell-type resolution^34^. To address this, patients were stratified by causal gene mutation (healthy donors n=18, *LMNA* n=11, *TTN* n=12, *RBM20* n=6, PVneg n=8) and ATP synthesis fluxes were evaluated across all five cell types as an index of overall energetic capacity (**Fig. 3A**). *LMNA*-mutant hearts exhibited mild cardiomyocyte ATP reduction (77% of healthy donors, FDR = 0.071) with more pronounced stromal suppression (endothelial 61%, FDR = 0.026; fibroblast 58%, FDR = 0.028; mural 60%, FDR = 0.010). *TTN*-mutant hearts retained relatively preserved cardiomyocyte metabolism (84%, FDR = 0.145). *RBM20*-mutant hearts displayed significant reductions in cardiomyocytes (70%, FDR = 0.044), endothelial cells (66%, FDR = 0.043), and mural cells (58%, FDR = 0.010). The PVneg group exhibited the most profound ATP depletion across all five cell types with cardiomyocytes at 64% (FDR = 0.050), endothelial cells at 47% (FDR = 0.015), fibroblasts at 43% (FDR = 0.009), mural cells at 46% (FDR = 0.004), and myeloid cells at 52% (FDR = 0.013) of healthy donor levels.

**Figure 3.**
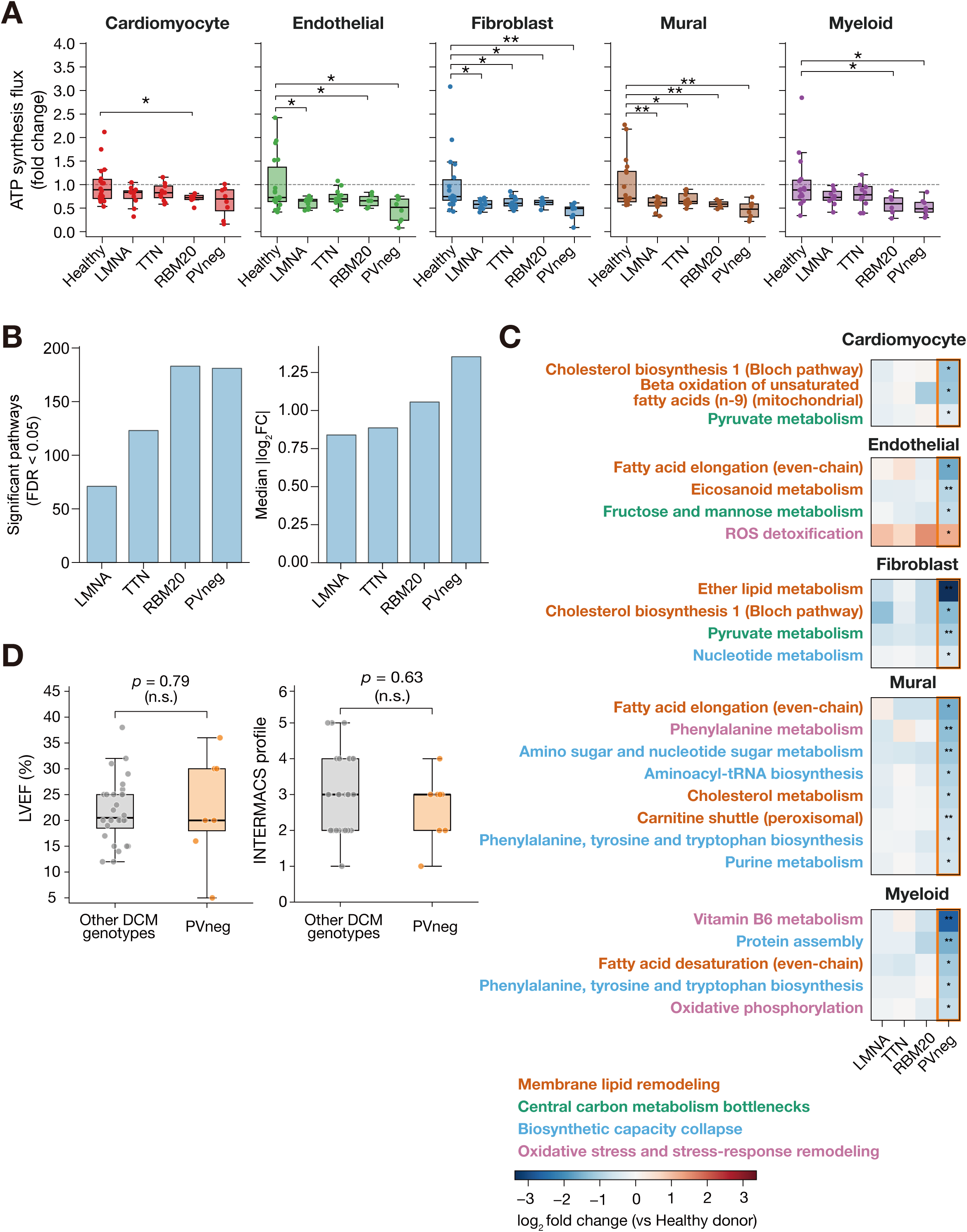
Genotype-dependent metabolic severity gradient in DCM. **(A)** ATP synthesis flux by genotype across five cell types. Each genotype (healthy donors n=18, *LMNA* n=11, *TTN* n=12, *RBM20* n=6, PVneg n=8) was compared to healthy donors. Welch’s *t*-test with Benjamini–Hochberg correction. \**p* < 0.05, \*\**p* < 0.01**. (B)** Genotype-dependent severity gradient quantified across all significantly altered pathway–cell type combinations (FDR < 0.05). Left: number of significantly altered pathways. Right: median |log_2_FC| of significant pathways. The median |log_2_FC| increased progressively across genotypes, while the number of altered pathways plateaued between *RBM20* and PVneg. **(C)** PVneg-specific metabolic pathways. Heatmap of 24 pathways significantly altered exclusively in PVneg (FDR < 0.05) while remaining non-significant (FDR > 0.1) in all other genotypes. Rows are grouped by cell type; columns represent genotypes. Asterisks denote FDR significance (\**p* < 0.05, \*\**p* < 0.01; PVneg vs healthy donors). **(D)** Clinical severity is comparable between PVneg and other DCM genotypes. Left: LVEF in PVneg (n=8) versus other DCM genotypes (*LMNA*, *TTN*, *RBM20*; n=29). Mann-Whitney U *p* = 0.79. Right: INTERMACS profile^37^. Mann-Whitney U *p* = 0.63. Both groups showed equivalent clinical severity, confirming that the observed metabolic differences are not confounded by differences in clinical severity.

To determine whether this hierarchy reflected a consistent gradient at the pathway level, we quantified the number of significantly altered pathways and the magnitude of their changes (FDR < 0.05) for each genotype across all cell type– pathway combinations (**Fig. 3B**). The median |log_2_FC| of significantly altered pathways increased progressively from *LMNA* (0.67) through *TTN* (0.71), *RBM20* (0.84), to PVneg (1.08). The number of altered pathways rose steeply from *LMNA* (71 pathways) to *TTN* (123) and reached a comparably high level in *RBM20* (183) and PVneg (181). This gradient is mechanistically coherent. Structural protein mutations in *LMNA* (nuclear envelope)^8^ and *TTN* (sarcomere)^7^ are expected to affect metabolism indirectly and modestly, whereas *RBM20* mutations perturb RNA splicing of metabolic genes^35,36^, producing intermediate disruption. The consistently severe metabolic impairment in PVneg hearts, despite the absence of identifiable structural gene mutations, suggests that metabolic dysfunction in this group represents a primary pathological process rather than a secondary consequence of structural defects.

The quantitative gradient, however, did not fully account for the metabolic phenotype of PVneg hearts. Examination of genotype-specific pathway alterations revealed 24 pathways significantly altered exclusively in the PVneg group (FDR < 0.05 in PVneg; FDR > 0.1 in all other genotypes; **Fig. 3C**), spanning all five cell types and converging on four mechanistic themes. First, widespread membrane lipid remodeling was evident, with cholesterol biosynthesis (Bloch pathway) suppressed in cardiomyocytes and fibroblasts, ether lipid synthesis reduced and fatty acid elongation diminished in endothelial and mural cells, and eicosanoid metabolism suppressed in endothelial cells, collectively indicating disruption of membrane composition and signaling lipid production. Second, central carbon metabolism was further impaired, with pyruvate metabolism reduced in cardiomyocytes and fibroblasts, revealing a bottleneck at the glycolysis–TCA cycle junction that extends beyond the common severity gradient. Third, biosynthetic capacity was broadly compromised, with aminoacyl-tRNA biosynthesis and purine metabolism suppressed in mural cells and nucleotide metabolism reduced in fibroblasts, indicating erosion of the fundamental machinery for protein and nucleic acid synthesis. Fourth, oxidative stress responses were augmented, with ROS detoxification upregulated in endothelial cells concurrent with reduced ether lipid (plasmalogen) synthesis, reflecting heightened oxidative stress in the PVneg stromal microenvironment and compensatory detoxification responses. Collectively, these four themes define PVneg DCM not as a more severe form of the common DCM metabolic phenotype but as a qualitatively distinct pathological entity.

Importantly, these genotype-dependent metabolic differences — both the severity gradient and the PVneg-specific signatures — were not confounded by clinical severity (**Fig. 3D**). Left ventricular ejection fraction (LVEF) did not differ between PVneg (median 20%) and other DCM genotypes (median 20.5%; Mann-Whitney U *p* = 0.79), nor did INTERMACS profile^37^ (median 3.0 vs. 3.0; *p* = 0.63), confirming that the metabolic gradient and PVneg-specific signatures reflect genotype-intrinsic programs independent of hemodynamic disease stage.

### Cross-cohort validation of metabolic alterations and genotype-specific patterns

To assess whether the metabolic remodeling identified in the Reichart cohort was reproducible, an identical pipeline was applied to two independent snRNA-seq cohorts (Chaffin^10^: 16 healthy donors, 8 DCM; Koenig^11^: 25 healthy donors, 12 DCM), and directional concordance of log_2_ fold change (DCM versus healthy donors) was computed across all three cohorts for the four cell types common to all datasets (cardiomyocyte, endothelial, fibroblast, and mural; **Table S3**). The overall three-way concordance was 52.0% (289/556 cell type–pathway combinations). Notably, core energy pathways, including oxidative phosphorylation, glycolysis, glycerolipid metabolism, and pyruvate metabolism, were consistently downregulated across all four cell types in all three cohorts (16/16 combinations negative), establishing pan-cellular energy suppression as the most robust and reproducible feature of DCM metabolism (**Fig. S2A**). Stromal cells exhibited higher concordance rates (fibroblast 61.2%, endothelial and mural 56.1%) than cardiomyocytes (34.5%), consistent with the greater directional consistency of stromal metabolic alterations observed in the discovery cohort.

Having established the reproducibility of pan-cohort metabolic patterns, we next assessed whether the genotype-specific features in the Reichart cohort were similarly reproducible. Using the Chaffin cohort (*TTN* n=4, PVneg n=4), in which the limited sample size precluded widespread FDR significance, ATP synthesis flux was nonetheless lower in PVneg than in *TTN*-mutant hearts across all five cell types (*TTN*: FC 0.76–0.89; PVneg: FC 0.60–0.69; **Fig. S2B**), reproducing the pattern of greater metabolic suppression in PVneg donors. To further test whether the 24 PVneg- specific pathways identified in the Reichart cohort reflected genuine PVneg biology rather than cohort-specific noise, we compared the log_2_ fold change direction in the Chaffin cohort for both PVneg and *TTN* (**Fig. S2C**). PVneg showed concordant direction in 21 of 24 pathways (88%), whereas *TTN* showed concordance in only 18 of 24 (75%), indicating that PVneg-specific metabolic signatures are not only reproducible in an independent cohort but are also specific to the PVneg genotype.

### Cross-disease analysis reveals shared ATP depletion and cell-type-dependent similarity across cardiomyopathies

To determine whether the metabolic remodeling observed in DCM represents a disease-specific phenomenon or a common feature of cardiomyopathies, the identical pipeline was applied to ACM (Reichart cohort, n=8) and HCM (Chaffin cohort, n=15). ATP synthesis flux was reduced across all cell types in both ACM and HCM (**Fig. 4A, B**), establishing that ATP depletion is a shared hallmark of cardiomyopathy regardless of etiology.

**Figure 4.**
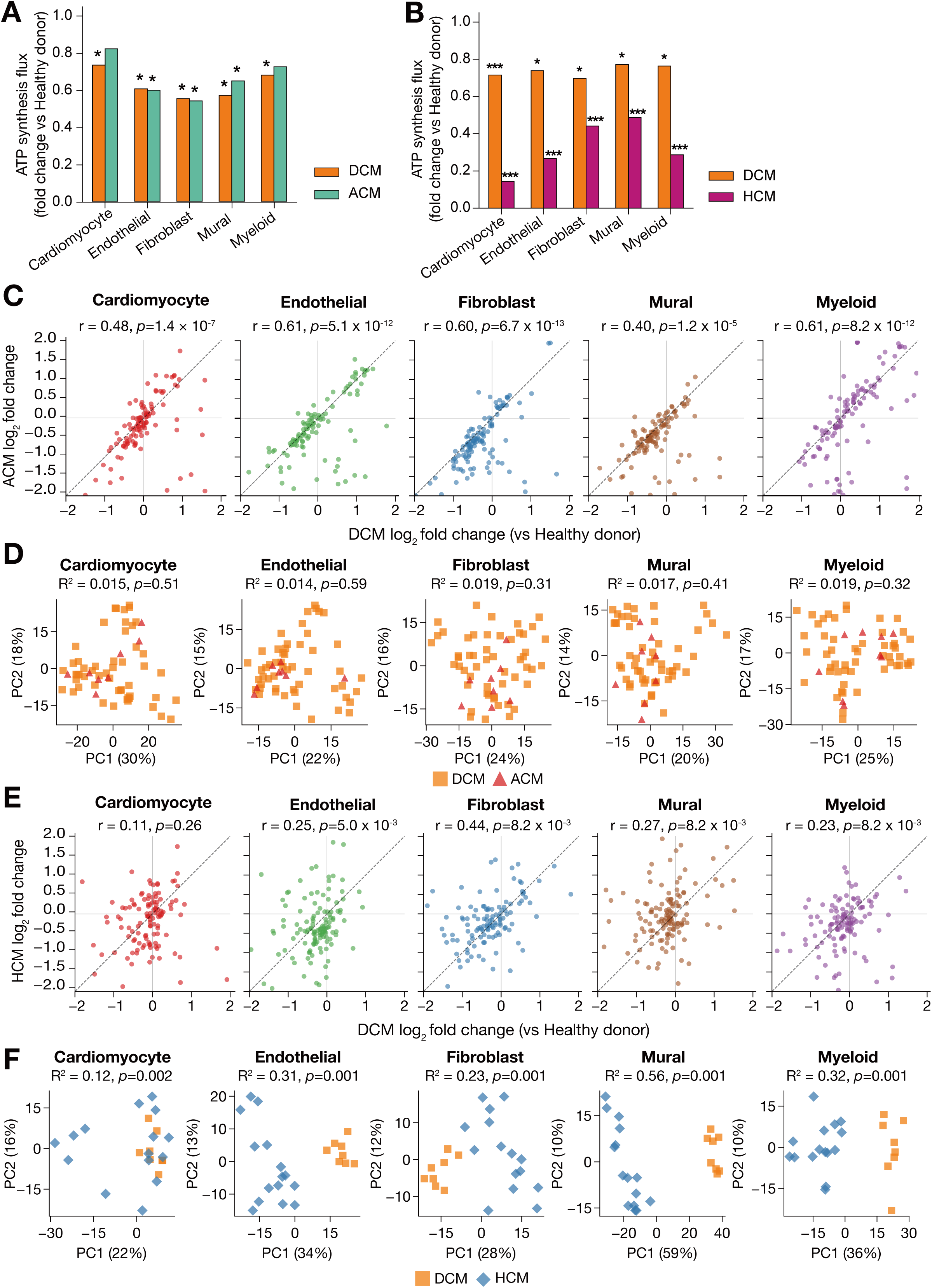
Cross-disease metabolic change in cardiomyopathy. **(A)** ATP synthesis flux fold change across five cell types in DCM (n=52) and ACM (n=8) relative to healthy donors (n=18) in the Reichart cohort^9^. Welch’s *t*-test with Benjamini–Hochberg correction. DCM showed significant ATP decline across all cell types (FC = 0.55–0.75, all FDR < 0.05); ACM showed a similar but attenuated pattern (FC = 0.59–0.81, 3/5 cell types significant). **(B)** ATP synthesis flux fold change in DCM (n=8) and HCM (n=15) relative to shared healthy donors (n=16) in the Chaffin cohort^10^. HCM exhibited more severe reductions (FC = 0.14–0.49, all FDR < 0.001) than DCM (FC = 0.70–0.77, all FDR < 0.05). **(C)** Scatter plots comparing pathway-level log_2_ fold changes (versus healthy donors) between DCM and ACM across 139 metabolic pathways for each of the five cell types. Each point represents one pathway. Pathways with |log_2_ fold change| ≥ 2 in either disease group were omitted from visualization. Dashed line indicates y = x. Pearson’s r and nominal *p* value are shown in each panel. DCM and ACM exhibited broadly similar pathway-level remodeling across the five cell types. **(D)** Principal component analysis (PCA) of donor-level pathway profiles for DCM and ACM in the Reichart cohort. For each donor and cell type, the input vector consisted of 139 pathway scores expressed as log_2_ (pathway score / healthy donor mean). Group separation was additionally quantified by permutational multivariate analysis of variance (PERMANOVA)^52^ in the original 139- dimensional feature space (Euclidean distance, 999 permutations), where R^2^ denotes the proportion of the total multivariate variance that is explained by the disease label. DCM and ACM donors broadly overlapped across the five cell types, with no statistically detectable separation in all five cell types (all R^2^ < 0.02, all *p* > 0.3). **(E)** Scatter plots comparing pathway-level log_2_ fold changes (versus healthy donors) between DCM and HCM across 139 metabolic pathways for each of the five cell types. Overall pathway-level similarity between DCM and HCM is lower than that observed between DCM and ACM, particularly in cardiomyocytes, indicating greater metabolic divergence in HCM. **(F)** PCA of donor-level pathway profiles for DCM and HCM in the Chaffin cohort, generated as in **(D)**. Fibroblasts showed the least clear separation in non-cardiomyocytes (PERMANOVA R^2^ = 0.227, *p* = 0.001).

We next compared metabolic pathway shifts within each cohort using all 139 metabolic pathways. In the Reichart cohort, DCM and ACM showed positive pathway-level correspondence across all five cell types (**Fig. 4C**) and broad overlap of donor-level profiles in principal component analysis (PCA) space (**Fig. 4D**), supporting convergent metabolic remodeling between these two disease groups. In the Chaffin cohort, DCM and HCM were less similar at the pathway level (**Fig. 4E**) and showed clear donor-level separation in several non-cardiomyocyte compartments (**Fig. 4F**), indicating greater subtype-dependent divergence in HCM relative to ACM. Across both comparisons, fibroblast profiles remained comparatively conserved, consistent with the disease-independent stromal metabolic suppression identified in DCM. Together, these results support shared ATP depletion across cardiomyopathies, close alignment between DCM and ACM, and fibroblast metabolic conservation as a recurrent feature across disease subtypes.

### Identification of transcription factor regulatory networks driving metabolic remodeling

The broad and coordinated nature of stromal metabolic suppression raises the question of whether a common transcriptional regulatory program underlies this remodeling. To elucidate the upstream drivers, transcription factor (TF) regulatory activity was analyzed across 11,554 metacells^38^ from 78 donors, identifying 128 regulons (TF–target gene sets) as the basis for subsequent analysis (see Methods). UMAP dimensionality reduction of metacell-level regulon activity scores (AUCell) revealed clear separation of the five major cell types (**Fig. 5A**). In contrast, healthy donor and DCM metacells were interspersed within each cell-type cluster (**Fig. 5B**), indicating that regulon activity is predominantly organized by cell lineage and that disease-associated changes occur within, rather than between, cell-type transcriptional programs.

**Figure 5.**
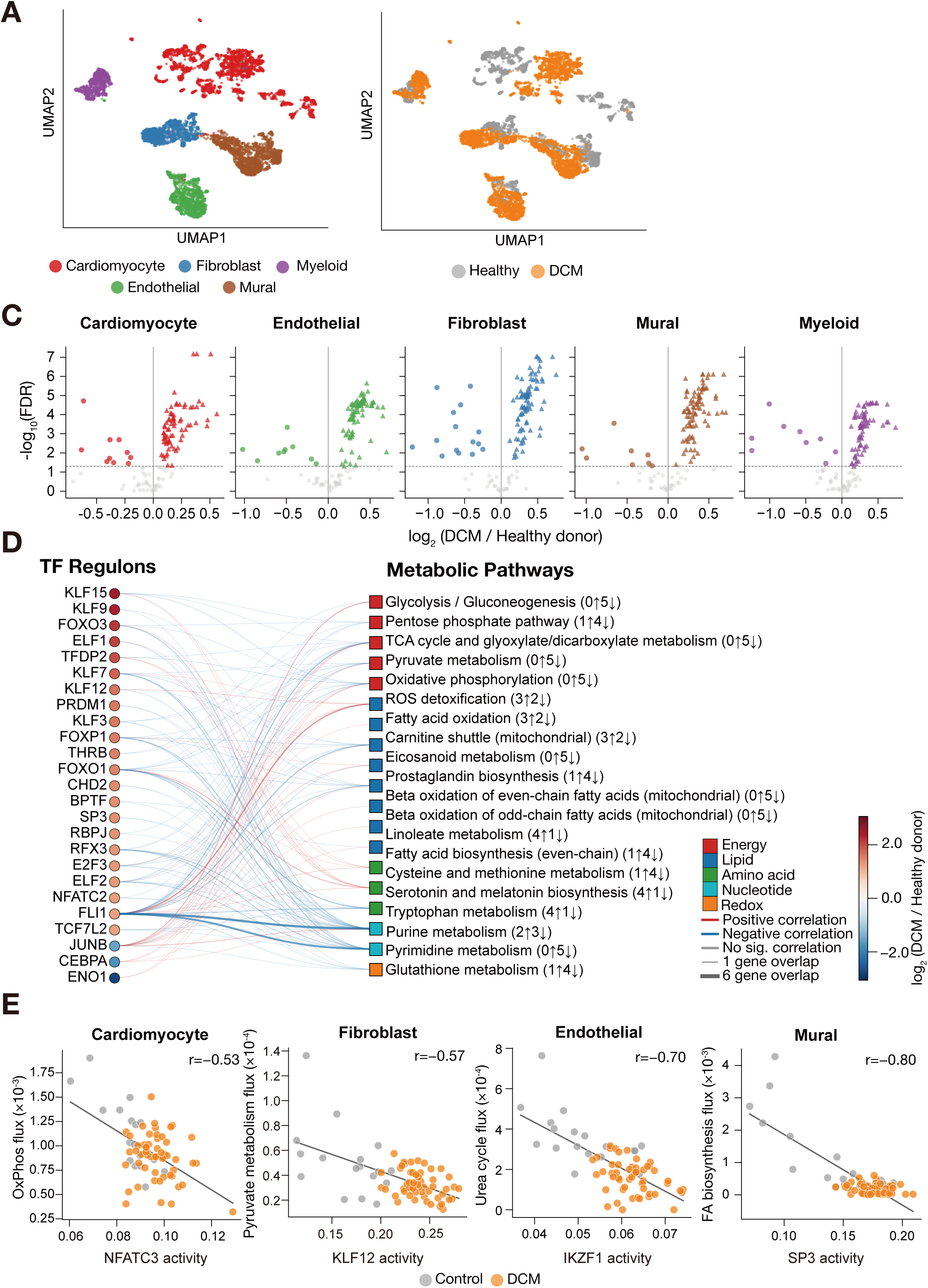
Transcription factor regulatory networks driving metabolic remodeling in DCM. **(A)** UMAP visualization of regulon activity profiles by cell type. AUCell scores for 128 regulons were computed on 11,554 metacells constructed per donor using SEACells^38^ from 78 donors. **(B)** Same UMAP as (A), colored by disease status. Healthy donor and DCM metacells were interspersed within each cell-type cluster, indicating that regulon activity variation is predominantly governed by cell-type identity. **(C)** Volcano plots of regulon activity changes (DCM versus healthy donor) for each cell type. Each point represents one regulon. Filled triangles indicate regulons significantly upregulated in DCM (FDR < 0.05, log_2_FC > 0); filled circles indicate regulons significantly downregulated (FDR < 0.05, log_2_FC < 0); gray points are non- significant. Dashed horizontal line marks FDR = 0.05. 454 of 640 cell type–regulon combinations (71%) were significantly altered. **(D)** Bipartite network linking transcription factor (TF) regulons to metabolic pathways through shared target genes (Human1^24^). Edge thickness is proportional to the number of overlapping genes. Numbers in parentheses after each pathway name indicate the number of cell types showing increased (↑) or decreased (↓) metabolic flux in DCM versus healthy donors. **(E)** Scatter plots of TF regulon activity versus metabolic pathway flux for four representative pairs (FDR < 0.05) spanning four cell types: Cardiomyocyte NFATC3 versus oxidative phosphorylation (Pearson’s r = −0.53), Fibroblast KLF12 versus pyruvate metabolism (r = −0.57), Endothelial IKZF1 versus urea cycle (r = −0.70), and Mural SP3 versus fatty acid biosynthesis (r = −0.80). The NFATC3–oxidative phosphorylation association is consistent with established calcineurin–NFAT signaling^41^; the remaining three represent novel associations.

Against this cell-type-structured background, DCM imposed widespread TF dysregulation. Cell-type-specific comparison of regulon activity between DCM and healthy donors revealed that 454 of 640 cell type–regulon combinations (71%) were significantly altered (FDR < 0.05; **Fig. 5C, Table S4**). Unlike the asymmetric metabolic flux alterations (**Fig. 2**), in which stromal pathways were preferentially suppressed, the number of significantly altered regulons was comparable across all five cell types (84–100), indicating that transcriptional dysregulation is a pan-cellular phenomenon that precedes or accompanies the cell-type-specific metabolic changes. To determine whether this transcriptional dysregulation is functionally coupled to the observed metabolic remodeling, we performed overlap analysis between regulon target genes and Human1^24^ metabolic pathway gene sets (**Fig. 5D**). Key metabolic pathways (oxidative phosphorylation, glycolysis, pentose phosphate pathway, glutathione metabolism, fatty acid oxidation, and TCA cycle) were connected to multiple FDR-significant regulons through shared target genes. The biological validity of these connections was supported by the identification of established cardiac metabolic regulators among the linked TFs, including KLF15, a critical transcriptional regulator of cardiac lipid metabolism whose expression is markedly reduced in human heart failure^39^, and FOXO1, which orchestrates cardiac substrate utilization and is a key driver of metabolic dysfunction in cardiomyopathy^40^.

To move beyond network topology and quantify the functional relationship between TF activity and metabolic flux, Pearson correlation analysis between TF regulon activity and metabolic pathway flux at the donor level identified 18,955 pairs with significant correlation (FDR < 0.05). Significant pairs were distributed across all five cell types: fibroblasts (5,505 pairs), mural cells (4,578 pairs), endothelial cells (3,909 pairs), myeloid cells (3,278 pairs), and cardiomyocytes (1,685 pairs), revealing pervasive TF–metabolic co-variation across the cardiac cellular landscape. These significant pairs encompassed both known and previously unreported TF–metabolic regulatory associations (**Fig. 5E**). The inverse correlation between NFATC3 activity and oxidative phosphorylation in cardiomyocytes (Pearson’s r = −0.53, FDR = 7.0 × 10^-5^) is consistent with prior evidence that calcineurin–NFAT signaling is accompanied by mitochondrial dysfunction during pathological cardiac hypertrophy^41^. In contrast, the inverse correlation between KLF12 activity and pyruvate metabolism flux in fibroblasts (r = −0.57, FDR = 8.3 × 10^-6^), the inverse correlation between the Ikaros family TF IKZF1 and urea cycle flux in endothelial cells (r = −0.70, FDR = 5.0 × 10^-9^), and the inverse correlation between SP3 activity and fatty acid biosynthesis in mural cells (r = −0.80, FDR = 8.9 × 10^-13^) represent novel TF–pathway associations in the cardiac context that have not been previously reported.

These results demonstrate a transcriptional regulatory architecture in which pan-cellular TF dysregulation is functionally coupled to metabolic flux remodeling across all cardiac cell types. Notably, regulon activity did not differ significantly among DCM genotypes (*LMNA*, *TTN*, *RBM20*, PVneg; **Fig. S3**), suggesting that transcriptional dysregulation is a genotype-independent common feature of DCM unlikely to account for the genotype-specific metabolic differences identified above.

### Machine learning classification reveals that metabolic pathway activity captures the molecular pathology of cardiomyopathy

To assess the informational content of cell-type-resolved metabolic pathway scores, metabolic pathway activity scores across all five cell types were used as features for machine learning classification and regression (5 cell types × 139 pathways; 695 features per donor, see Methods). Disease-state classification of healthy donors versus DCM achieved near-perfect accuracy, with all models reaching AUC ≥ 0.90 (**Fig. 6A, Table S5**). LASSO coefficient analysis identified fibroblast glycerolipid metabolism (coefficient = −0.89) as the top discriminator, followed by mural cell eicosanoid metabolism and fatty acid biosynthesis (odd-chain) (**Fig. 6B, Table S6**). The multi-cell-type composition of the top features directly reinforces the conclusion that DCM-associated metabolic remodeling is a pan-cellular rather than cardiomyocyte-restricted phenomenon.

**Figure 6.**
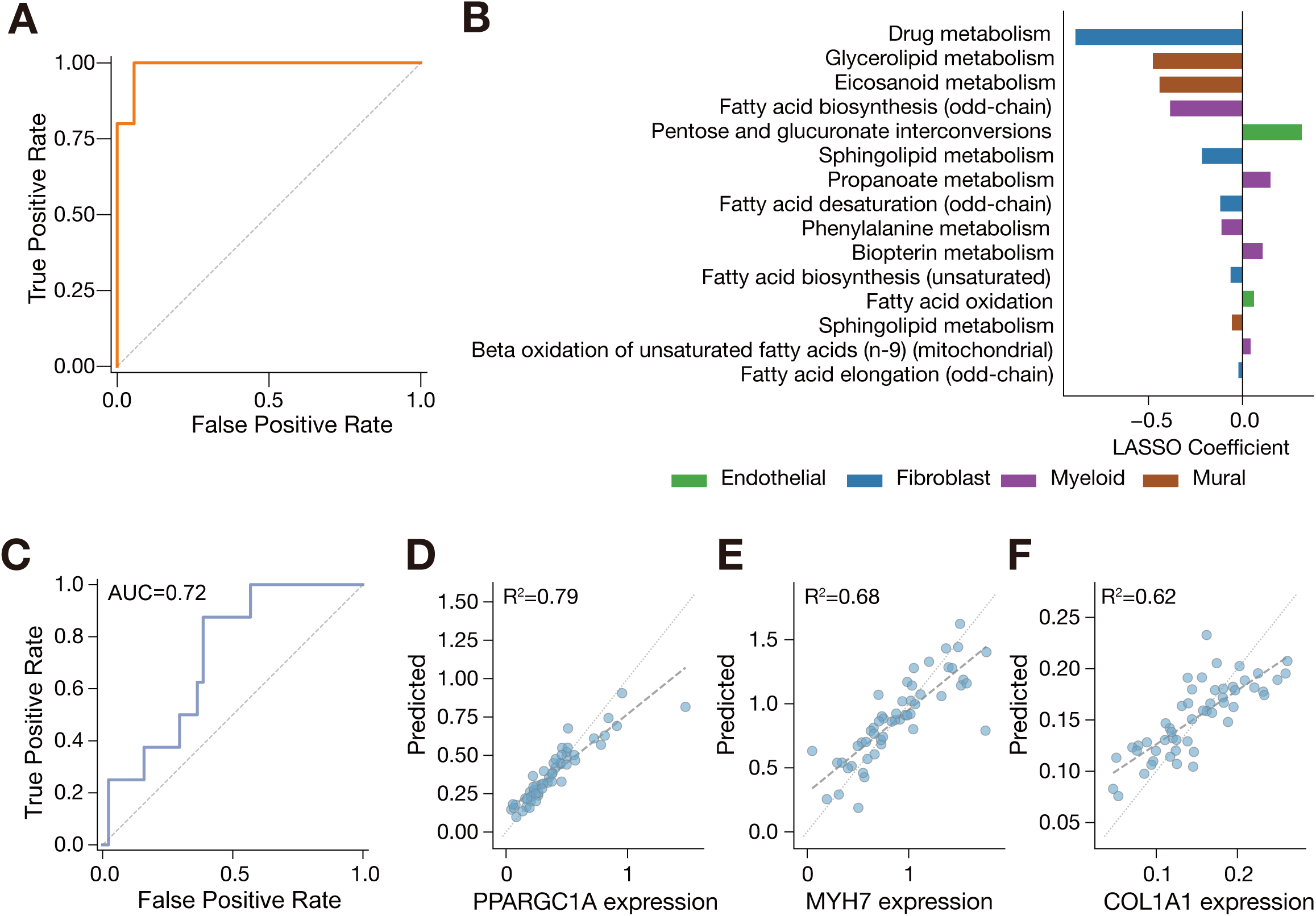
Machine learning classification reveals that metabolic pathway scores capture the molecular pathology of cardiomyopathy. **(A)** ROC curve for healthy donors versus DCM classification using LASSO logistic regression with Leave-One-Out Cross-Validation (LOOCV; healthy donors n=18, DCM n=52; AUC=0.980). All seven models achieved AUC ≥ 0.90, with SVM linear reaching AUC = 0.999. **(B)** LASSO coefficient bar plot showing the top 15 non-zero features for healthy donors versus DCM classification. Features are colored by cell type. Fibroblast glycerolipid metabolism exhibited the largest absolute coefficient (−0.89). **(C)** ROC curve for PVneg (n=8) versus non-PVneg DCM (n=44) classification using Random Forest (AUC = 0.72) with LOOCV. **(D–F)** Scatter plots of observed versus predicted marker gene expression from metabolic pathway scores (LOOCV, n=52 DCM donors). **(D)** *PPARGC1A*^42^ (R^2^ = 0.79). **(E)** *MYH7*^43^ (R^2^ = 0.68). **(F)** *COL1A1*^44^ (R^2^ = 0.62).

Having established that metabolic pathway scores distinguish disease from health, we asked whether they could further resolve genotypic heterogeneity within DCM. Classification of PVneg versus non-PVneg DCM achieved moderate separation (Random Forest AUC = 0.72; **Fig. 6C, Table S5**), consistent with the quantitative and qualitative genotype-dependent metabolic differences identified in earlier analyses (**Fig. 3**). To assess whether metabolic pathway activity scores reflect not only disease state but also the degree of molecular reprogramming, regression analysis was performed to predict individual marker gene expression from metabolic pathway scores in DCM donors. Prediction accuracy was high for markers of mitochondrial biogenesis (*PPARGC1A*^42^, R^2^ = 0.79), cardiac contraction (*MYH7*^43^, R^2^ = 0.68), and fibrosis (*COL1A1*^44^, R^2^ = 0.62) (**Fig. 6D-F**; **Table S7**), demonstrating that metabolic pathway scores quantitatively capture the extent of transcriptional reprogramming across these cardinal pathological processes. Notably, clinical indices of heart failure (LVEF, LVIDd, eGFR, BNP) were poorly predicted from the same pathway scores (R^2^ ≤ 0.07; **Fig. S4**; **Table S8**), indicating that the metabolic remodeling identified here reflects molecular pathology intrinsic to cardiomyopathy rather than the hemodynamic consequences of pump failure.

## Discussion

In this study, we revealed that cardiomyopathy-associated metabolic remodeling is a pan-cellular process with pronounced stromal reprogramming (**Fig. 7**). ATP synthesis was reduced across all five major cardiac cell types, with the strongest impairment in fibroblast, mural, and endothelial cells, whereas myeloid cells exhibited relative activation of stress-associated metabolic pathways. Within DCM, metabolic impairment followed a genotype-dependent severity gradient, and pathogenic variant-negative hearts harbored a distinct set of metabolic alterations independent of clinical severity. Cross-disease analyses further indicated shared ATP depletion across cardiomyopathy subtypes and comparatively conserved fibroblast profiles across disease comparisons.

**Figure 7.**
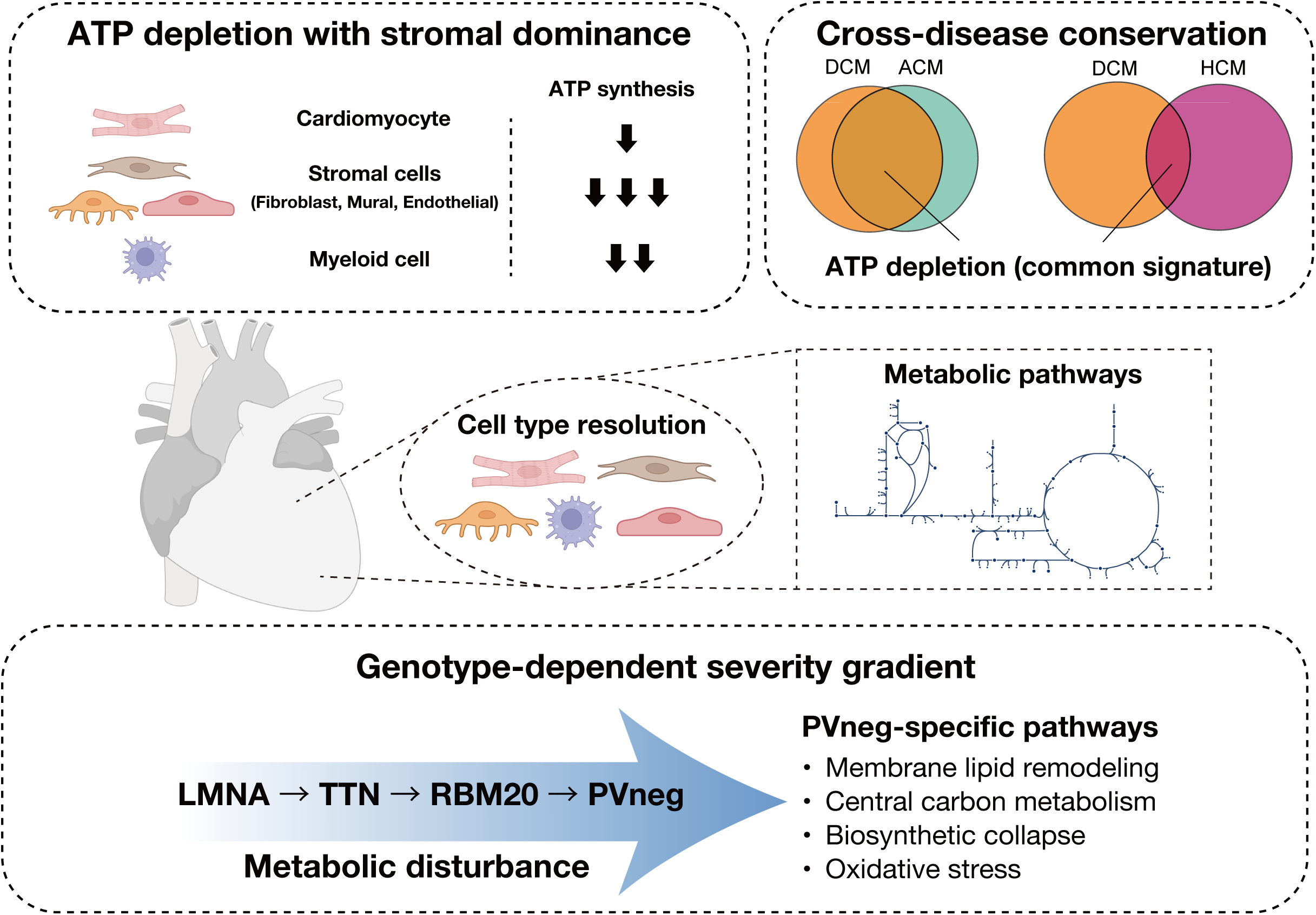
Schematic summary of cell-type-resolved metabolic remodeling in cardiomyopathy. Single-nucleus transcriptomes from human hearts were resolved into five major cardiac cell types (cardiomyocytes, endothelial cells, fibroblasts, mural cells, and myeloid cells) and used to reconstruct cell-type–specific metabolic networks. **(Top left)** ATP synthesis flux is reduced across all five major cardiac cell types in dilated cardiomyopathy (DCM), with the most pronounced impairment observed in stromal populations (fibroblasts, mural cells, and endothelial cells), and a comparatively milder reduction in cardiomyocytes and myeloid cells. **(Top right)** Cross-disease comparison reveals that ATP depletion constitutes a shared metabolic signature across cardiomyopathy subtypes, with DCM and arrhythmogenic cardiomyopathy (ACM) exhibiting more extensive overlap than DCM and hypertrophic cardiomyopathy (HCM). **(Bottom)** Within DCM, metabolic disturbance follows a genotype-dependent severity gradient. Pathogenic variant–negative (PVneg) hearts further harbor a distinct set of metabolic alterations, including membrane lipid remodeling, central carbon metabolism dysregulation, biosynthetic collapse, and oxidative stress, that are independent of clinical severity indices such as LVEF and INTERMACS profile. Together, these findings position cardiomyopathy as a pan-cellular metabolic disease with stromal-dominant ATP impairment, a genotype-graded severity axis, and a distinct PVneg metabolic phenotype.

Previous bulk-tissue metabolomic studies have documented energy starvation^12^, depletion of fatty acid intermediates^13^, and a substrate shift toward ketone body utilization in failing hearts^14^. Computational transcriptomic approaches have further identified disruptions in intercellular metabolic signaling^45^ and perturbations in arachidonic acid metabolism across cardiomyopathies^46^. However, these findings are based on tissue-level averages and cannot resolve the cell-type distribution of metabolic dysfunction. The present study demonstrates at cell-type resolution that the most profound ATP production deficits in DCM reside in stromal cells, with fibroblasts (FC = 0.55), mural cells (FC = 0.58), and endothelial cells (FC = 0.61), all showing greater energetic impairment than cardiomyocytes (FC = 0.75; **Fig. 2A**). This finding extends the transcriptional-level observations of dramatic fibroblast diversification^11^ and the emergence of disease-specific activated fibroblast populations^10^ to the level of metabolic flux, and establishes stromal energy failure as a quantitatively dominant feature of the failing heart. In contrast, myeloid cells exhibited marked upregulation of tryptophan metabolism and steroid metabolism, revealing a compartmentalized response in which stromal energy failure coexists with immune metabolic activation (**Fig. 2D**).

DCM-associated metabolic impairment was not uniform across genotypes but followed a severity gradient from *LMNA* to *TTN*, *RBM20*, and PVneg (**Fig. 3A, B**). Structural protein mutations in *LMNA* and *TTN* exerted indirect and modest effects on metabolism, whereas PVneg hearts exhibited the most profound ATP depletion across all five cell types. Beyond this quantitative gradient, 24 metabolic pathways were exclusively altered in PVneg hearts, converging on membrane lipid remodeling, central carbon metabolism bottlenecks, biosynthetic capacity collapse, and augmented oxidative stress (**Fig. 3C**). The independence of these signatures from clinical severity measures (**Fig. 3D**) indicates that they reflect genotype-intrinsic metabolic programs rather than consequences of more advanced disease, positioning PVneg DCM as a pathophysiologically distinct entity in which primary metabolic dysfunction may be a disease driver rather than a secondary manifestation. The specific pathway alterations identified, including cholesterol biosynthesis, ether lipid synthesis, and aminoacyl-tRNA biosynthesis, represent candidate therapeutic targets for genotype-informed intervention, warranting validation in cellular and animal models of PVneg DCM.

The cross-disease comparison refines the interpretation of shared versus subtype-dependent remodeling. ATP depletion was observed not only in DCM but also in ACM and HCM (**Fig. 4A, B**), indicating that energetic failure is a common feature of cardiomyopathy. At the pathway level, DCM and ACM were more closely aligned than DCM and HCM, suggesting that dilated cardiomyopathies of different etiologies converge on related metabolic programs (**Fig. 4C, E**). Donor-level analyses further supported this interpretation: DCM and ACM were not detectably separated (**Fig. 4D**), whereas DCM and HCM showed clearer separation in several non-cardiomyocyte compartments, strongest in mural cells and weakest in fibroblasts (**Fig. 4F**). Cardiomyocyte profiles did not show clear separation between DCM and HCM (**Fig. 4F**). However, cardiomyocytes already display only modest pathway-level changes relative to healthy donors in DCM (**Fig. 2C**), limiting the strength of conclusions regarding disease-specific cardiomyocyte remodeling. A more defensible interpretation is that fibroblast metabolic reprogramming represents a conserved stromal response across cardiomyopathies, with additional subtype-dependent remodeling superimposed in other cellular compartments.

From a methodological standpoint, this study established a metabolic flux inference framework suited for non-proliferative tissues by adopting ATP synthesis maximization as the objective function. Conventional biomass maximization models underestimated cardiomyocyte energy metabolism pathways and equalized flux allocation across cell types, whereas the ATP maximization framework recapitulated physiologically valid cell-type-specific profiles (**Fig. 1B, C; Fig. S1**). The principle of tissue-specific objective function selection is broadly applicable, and the present framework may be extended to metabolic analysis of other non-proliferative organs such as the brain and skeletal muscle where ATP homeostasis similarly dominates over biomass accumulation.

This study has several limitations. First, inferred fluxes are predictions based on transcript abundance and stoichiometric models rather than direct measurements of enzymatic activity or metabolite concentrations. However, single-cell resolution quantification of intracellular metabolic fluxes in primary human tissue remains technically infeasible. Future advances in single-cell and spatial metabolomics may enable direct validation of cell-type-resolved flux estimates. Second, the ACM (n=8) and HCM (n=15) cohorts were limited in size, and the HCM data were derived from a different cohort (Chaffin cohort), precluding formal exclusion of cohort effects. Third, the analysis is cross-sectional, and whether metabolic dysfunction precedes, accompanies, or follows structural deterioration in cardiomyopathy requires longitudinal or interventional studies in appropriate animal models.

In conclusion, cell-type-resolved metabolic flux analysis revealed pan-cellular energetic impairment with pronounced stromal vulnerability, shared ATP depletion across cardiomyopathy subtypes, and a distinct metabolic program in pathogenic variant-negative DCM. These findings establish conserved stromal vulnerabilities as candidate therapeutic targets and demonstrate that tissue-specific flux modeling of transcriptomic atlases offers a resolution of human organ metabolism not achievable by bulk-tissue approaches.

## Methods

### Patient cohorts and snRNA-seq datasets

Three independent snRNA-seq datasets were used to analyze metabolic remodeling in cardiomyopathy.

#### Reichart cohort

A large-scale snRNA-seq dataset generated by Reichart et al. ^9^ and publicly available through the CELLxGENE portal served as the primary analysis cohort. The cohort comprised 79 donors: 52 dilated cardiomyopathy (DCM), 8 arrhythmogenic cardiomyopathy (ACM), 1 restrictive cardiomyopathy/left ventricular non-compaction (RCM/LVNC), and 18 healthy donors. Because the RCM/LVNC group consisted of a single donor, this case was excluded from subsequent analyses, resulting in a total of 78 donors included in the study. Cell-type annotations and preprocessing followed the original study pipeline.

#### Chaffin cohort (validation 1)

A DCM snRNA-seq dataset (24 donors: 16 healthy donors, 8 DCM) was obtained from the Single Cell Portal database^10^. Cell-type annotations were derived from clustering after CPM normalization, log1p transformation, regression of library size and mitochondrial fraction on highly variable genes, and Harmony batch correction^47^. This dataset additionally included 15 HCM patients.

#### Koenig cohort (validation 2)

A mixed scRNA-seq/snRNA-seq dataset (37 donors: 25 healthy donors, 12 DCM, with the snRNA-seq subset exclusively) was obtained from Koenig et al.^11^ (GEO: GSE183852). Cell-type annotations were generated de novo using the same preprocessing and clustering pipeline as the Reichart cohort.

Lymphocyte and myeloid cell annotations were not available in this dataset, and analyses were therefore restricted to four cell types (cardiomyocyte, endothelial, fibroblast, mural).

The Reichart cohort contained 869,449 nuclei annotated into ten cell types. We retained five major cell types (cardiomyocyte, fibroblast, endothelial, mural, myeloid) with sufficient cell numbers for reliable pseudobulk profiling. Cross-cohort validation used the four cell types common to all three cohorts (cardiomyocyte, endothelial, fibroblast, mural).

### Cell-type-resolved cardiac metabolic flux inference

Building on the METAFlux^20^, a community flux balance analysis framework, which jointly models multiple cell types within a tissue by incorporating cell-type fractions as stoichiometric constraints, we estimated metabolic fluxes using the Human1^24^, a genome-scale metabolic model (13,082 reactions, 3,625 genes), with cardiac- specific modifications as detailed below.

### Pseudobulk expression profiles

Donor-level pseudobulk expression profiles were generated as follows. For each donor, raw gene counts were aggregated across all cells within each cell type using decoupler^48^, yielding one expression vector per donor–cell type combination. Counts were then normalized to counts per ten thousand and log-transformed (log1p) using scanpy^49^.

### Gene filtering and exclusion of mitochondrial DNA-encoded genes

To address zero-inflation inherent to snRNA-seq data, gene filtering was performed in two steps. First, for each cell type independently, only genes detected in all donors (pseudobulk expression greater than zero) were retained as flux constraints. In the Reichart cohort, the number of retained genes varied by cell type (approximately 15,000 for cardiomyocytes and 14,000 for endothelial cells), ensuring that only robustly measured genes served as metabolic flux constraints for each cell type. Second, because snRNA-seq captures only nuclear RNA, transcripts encoded by the mitochondrial genome (e.g. MT-ND1-6, MT-CYB, MT-CO1-3, MT-ATP6, MT-ATP8) reflect artifacts from ambient RNA or damaged cells rather than true mitochondrial transcriptional activity^50^ and were therefore excluded from all gene-protein-reaction (GPR) associations. Reactions in which the removed gene was the sole subunit were assigned a score of 1 (unconstrained), ensuring that fluxes through these reactions were governed exclusively by nuclear-encoded gene expression.

### Metabolic reaction activity scores

Metabolic Reaction Activity Scores (MRAS) were computed from filtered expression data using gene-protein-reaction (GPR) rules as implemented in METAFlux^20^. For enzyme complexes (AND relationships), the score was determined by the minimum expression value across subunit genes, divided by the number of reactions involving that gene. For reactions catalyzed by isoenzymes (OR relationships), the expression values of all isoenzyme genes were summed, reflecting the assumption that isoenzymes contribute additively to the total reaction capacity. MRAS values were max-normalized to a 0–1 range within each donor-cell type combination.

### Objective function and flux balance analysis

Rather than the conventional biomass maximization objective, we maximized ATP synthesis flux (reaction HMR_6916 in Human1^24^) to reflect the bioenergetic demands of non-proliferative cardiac tissue. The objective function was defined as

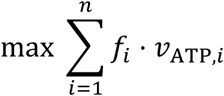

where 𝑓_𝑖_is the cell-type fraction of cell type 𝑖 and 𝑣_ATP,𝑖_is the corresponding ATP synthesis flux. The optimization was subject to four constraints: (1) steady-state mass balance; (2) flux bounds derived from MRAS-derived reaction directionality; (3) medium constraints defined by human blood metabolite composition (METAFlux default); and (4) cell-type fractions empirically derived from the observed cell-type proportions of each donor. This maximization problem was solved as a quadratic programming:

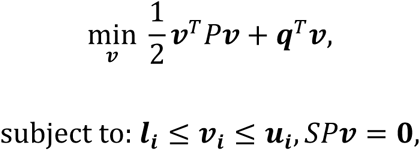

where 𝑃 is a diagonal weighting matrix whose entries are the cell-type fractions 𝑓_𝑖_ replicated over the 13,082 reactions of each cell type (with unit weights for the 1,648 external medium reactions). 𝒒 is a sparse linear coefficient vector with 𝑞_ATP,𝑖_ = − 10^4^𝑓_𝑖_ and 𝑞_otherwise_ = 0 , so that the linear term maximizes Σ𝑓_𝑖_𝑣_ATP,𝑖_ while the quadratic term 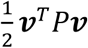 provides a cell-type-weighted L2 regularization on the flux vector 𝒗 . Flux bounds 𝒍_𝒊_ and 𝒖_𝒊_ were derived from the MRAS-based reaction directionality (upper bound = MRAS; lower bound = −MRAS for reversible reactions, 0 for irreversible reactions), with 𝒍_𝒊_ = −1 for exchange reactions of metabolites present in the human blood medium and 𝒍_𝒊_ = 0 otherwise, for each cell type 𝑖 . 𝑆 is the stoichiometric matrix. The QP was solved using the OSQP solver^51^ with a maximum of 1,000,000 iterations.

### Pathway activity analysis Pathway activity scores

Pathway activity scores were calculated for each donor for 139 metabolic pathways (subsystems) defined in Human1^24^, excluding transport, exchange, and pool reactions. For each donor and cell type, the pathway activity score was computed as the mean absolute flux across all reactions in the subsystem:

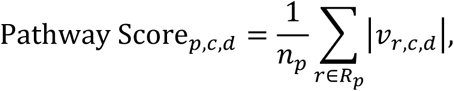

where 𝑝 denotes the pathway, 𝑐 the cell type, 𝑑 the donor, 𝑅_𝑝_ the set of reactions in subsystem 𝑝, 𝑛_𝑝_ the number of reactions, and 𝑣_𝑟,𝑐,𝑑_ the flux value through reaction 𝑟.

### Metabolic pathway activity analysis and physiological validation

For each pathway, mean absolute flux was computed per cell type across healthy donors (n=18), and Z-score standardization was applied across the five cell types:

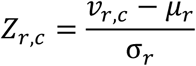

where 𝜇_𝑟_and 𝜎_𝑟_are the mean and standard deviation across all cell types, respectively. This standardization identifies which cell type preferentially utilizes each metabolic pathway relative to other cell types.

To validate the physiological relevance of predicted fluxes, 15 representative pathways were selected from five biological categories (cardiac energy, endothelial/vascular, fibroblast/ECM, mural/lipid, and myeloid/immune) based on published literature supporting the biological relevance of each pathway to its assigned cell type. Z-scores for all 139 Human1 pathways are provided in the **Supplementary Table 2**.

### Dimensionality reduction and cell-type clustering of metabolic profiles

To visualize cell-type-specific metabolic identities, dimensionality reduction was performed on pathway activity scores from healthy donors (n=18). For each donor– cell type combination, pathway activity scores were computed for 139 Human1 pathways and standardized to zero mean and unit variance per pathway. Principal component analysis (PCA) was applied to retain 50 components, followed by Uniform Manifold Approximation and Projection (UMAP^25^; n_neighbors=15, min_dist=0.1, random_state=42) to generate a two-dimensional embedding.

### Cell-type-specific pathway comparison

For each cell type–pathway combination, pathway activity differences between DCM and healthy donors were evaluated using Welch’s *t*-test (two-sided, unequal variance). Benjamini-Hochberg FDR correction was applied across all cell type–pathway combinations, with FDR < 0.05 as the significance threshold. Only combinations with pathway activity measured in at least three donors in both groups were included; combinations with zero variance were assigned *p* = 1.0.

### Genotype-stratified pathway analysis

To evaluate genotype-specific metabolic remodeling, DCM patients were stratified by causal gene mutation into four groups: *LMNA* (n=11)^8^, *TTN* (n=12)^7^, *RBM20* (n=6)^35^, and PVneg (pathogenic variant-negative, n=8). Healthy donors (n=18) served as the reference group. Donors with other rare genotypes were excluded from genotype-stratified analyses due to limited numbers. For each genotype group, pathway activity scores were computed per cell type as described above, and Welch’s *t*-test was performed for each cell type–pathway combination comparing each genotype group to the healthy donor group. Benjamini-Hochberg FDR correction was applied independently within each genotype, with FDR < 0.05 as the significance threshold. PVneg-specific pathways were defined as those reaching significance in the PVneg group (FDR < 0.05) while remaining non-significant in all other genotypes (FDR > 0.1 in *LMNA*, *TTN*, and *RBM20*).

### Cross-cohort concordance analysis

To assess the reproducibility of metabolic pathway alterations, an identical pipeline was applied to each of the three cohorts (Reichart, Chaffin, and Koenig) independently, and log_2_ fold changes and Welch’s *t*-tests for DCM versus healthy donors were computed separately for each cohort. For the four cell types shared across all three cohorts (cardiomyocyte, endothelial, fibroblast, and mural), concordance rate was defined as the proportion of cell type–pathway combinations in which the direction of log_2_ fold change (positive or negative) was consistent across all three cohorts. Pathways exhibiting the same direction of change in all three cohorts were designated as 3-way concordant and reported as reproducible metabolic alterations.

### Cross-disease pathway similarity and donor-level PCA analysis

To assess whether metabolic remodeling is shared across cardiomyopathy subtypes while minimizing cohort effects, cross-disease comparisons were performed within cohort: DCM versus ACM in the Reichart cohort and DCM versus HCM in the Chaffin cohort. For each disease group and cell type, pathway activity scores were calculated for all 139 Human1 pathways as described above and expressed as log_2_ fold change relative to the mean of healthy donors from the same cohort. Pathway-level scatter plots were generated by comparing log_2_ fold changes (versus healthy donors) across paired disease groups for each cell type. For visualization only, pathways with |log_2_ fold change| ≥ 2 in either disease group were omitted. Pearson correlation coefficients were used as descriptive measures of pathway-level similarity. For donor-level analyses, each disease donor was represented by a 139-dimensional vector defined as log_2_ (pathway score / healthy-donor mean) for the corresponding cohort and cell type. Principal component analysis (PCA) was performed independently for each cell type and disease comparison. Where indicated, donor-level separation was additionally summarized using permutational multivariate analysis of variance (PERMANOVA)^52^ on Euclidean distance matrices with 999 permutations using scikit-bio^53^ (v0.7.2). R^2^ denotes the proportion of the total multivariate variance in the original 139-dimensional feature space that is explained by the disease label.

### Transcription factor activity analysis and metabolic regulatory network Metacell construction

To reduce technical noise while preserving inter-donor biological variation, metacells were constructed independently for each of the 78 donors. Raw count data were extracted for each donor and preprocessed following the preprocessing method of the original Reichart cohort: CPM normalization (target sum of 1 × 10⁴ counts), log1p transformation, highly variable gene selection, regression of library size and mitochondrial fraction, scaling (clipped to a maximum value of 10), PCA (40 components, using the ARPACK solver), and neighbor graph construction (15 nearest neighbors). SEACells^38^ (v0.3.3) was applied to the preprocessed data to generate metacells, with the target number set to each donor’s cell count divided by 75 (minimum 10). Raw counts were summed within each metacell, and cell-type annotations were assigned by majority vote. Metacells from all 78 donors were concatenated and subjected to CPM normalization and log1p transformation, yielding a final expression matrix of 11,554 metacells × 33,145 genes.

### Gene regulatory network inference

Gene regulatory networks were inferred using the pySCENIC (v0.12.1)^23^, GRNBoost2– cisTarget–AUCell pipeline. The log-normalized metacell expression matrix and a curated list of 1,892 human transcription factors (from the pySCENIC human TF reference, hg38) were provided as input to the GRNBoost2 algorithm (arboreto^54^ v0.1.6; gradient boosting, random seed = 42) to compute TF–target gene adjacency scores. Co-expression modules were then constructed from the adjacencies within the pySCENIC pipeline and subjected to cisTarget motif enrichment pruning using two ranking databases for the hg38 genome (TSS ± 10 kbp and TSS +500 bp/−100 bp, version 10, Aerts Lab) and the corresponding motif annotation table (version 10, clustered, HGNC symbols). Pruning retained 128 regulons (TF–target gene sets) supported by motif evidence.

### AUCell activity scoring

Regulon activity was quantified using AUCell (Area Under the Recovery Curve) scores. The log-normalized metacell expression matrix (11,554 metacells) was used as input and the AUCell algorithm was applied with an AUC threshold of 0.05, corresponding to the top 5% of the gene expression ranking. AUCell measures the enrichment of regulon target genes among the top-ranked expressed genes of each cell, yielding a score between 0 and 1. Metacells were processed in chunks of 50,000 for memory efficiency.

### Donor-level aggregation

Metacell-level AUCell scores were aggregated to the donor level by averaging scores across all metacells within each donor–cell type combination. Analysis was restricted to the five major cell types (cardiomyocyte, endothelial, fibroblast, mural, myeloid).

### DCM versus healthy donor regulon activity comparison

For each cell type–regulon combination, donor-level AUCell scores were compared between healthy donors and DCM using two-sided Mann-Whitney U tests. Benjamini-Hochberg FDR correction was applied independently within each cell type, with FDR < 0.05 as the significance threshold.

### TF regulon–metabolic pathway overlap analysis

To assess functional coupling between TF regulons and metabolic pathways, we computed the overlap between each regulon’s target gene set and the constituent genes of Human1 metabolic pathways (extracted from gene-protein-reaction associations). TF–pathway pairs with one or more overlapping genes were defined as regulatory connections. For the bipartite network (**Fig. 5D**), the top 25 regulons were selected from those with ≥ 2 overlapping genes with key metabolic pathways, mean AUCell score ≥ 0.01 across all five cell types, and significant differential activity between healthy donor and DCM (FDR < 0.05), ranked by absolute log_2_ fold change.

### TF activity–metabolic flux correlation analysis

To evaluate co-variation between TF regulon activity and metabolic pathway flux, Pearson correlation analysis was performed at the donor level. For each cell type, correlation coefficients were computed for all regulon–pathway combinations with at least five common donors. Regulons or pathways with zero variance were excluded. Benjamini-Hochberg FDR correction was applied across all tests, with FDR < 0.05 as the significance threshold.

### Inter-genotype regulon activity comparison

To assess whether regulon activity differed among DCM genotypes (LMNA, TTN, RBM20, PVneg), Kruskal-Wallis tests were performed for each cell type–regulon combination. Benjamini-Hochberg FDR correction was applied across all tests, with FDR < 0.05 as the significance threshold. For the heatmap (**Fig. S3**), the same top 25 regulons selected for bipartite network (**Fig. 5D**) were used. Heatmap values represent log_2_ fold change of mean AUCell scores for each genotype relative to healthy donor.

### Machine learning classification and regression Feature construction

For each donor, pathway activity scores across five cell types (cardiomyocyte, endothelial, fibroblast, mural, and myeloid) and 139 Human1 pathways were computed, yielding 695 features.

### Disease-state classification

Seven classification models (LASSO, Ridge, ElasticNet, Random Forest, SVM linear, SVM RBF, LightGBM) were benchmarked using Leave-One-Out Cross-Validation (LOOCV) for healthy donors (n=18) versus DCM (n=52). ROC-AUC was computed for each model. Feature importance was assessed using non-zero LASSO coefficients, with the top 15 features reported.

### Pathogenic variant-negative classification

Classification of PVneg (n=8) versus non-PVneg DCM (n=44) was performed using three classification models (LASSO, Random Forest, LightGBM). Linear/kernel models were excluded due to LOOCV instability under extreme class imbalance (8/52). LOOCV-derived prediction probabilities were used to compute ROC-AUC.

### Marker gene expression prediction

To assess whether metabolic pathway scores capture transcriptional reprogramming, regression analysis was performed to predict individual gene expression from pathway scores in DCM donors (n=52, LOOCV). For each gene, the best-performing model (LASSO, Ridge, or LightGBM) was selected based on R^2^. Target genes included markers of mitochondrial biogenesis (*PPARGC1A*), cardiac contraction (*MYH7*, *MYH6*, *PLN*, *ATP2A2*), fibrosis (*COL1A1*, *COL3A1*, *POSTN*), and heart failure (*ANKRD1*, *NPPB*, *NPPA*).

### Clinical index prediction

To test whether metabolic remodeling correlates with clinical severity, regression analysis was performed to predict clinical indices (LVEF, LVIDd, eGFR, and BNP) from pathway scores in DCM donors. Four regression models (LASSO, Ridge, Random Forest, LightGBM) were evaluated using LOOCV. Outliers were defined as values beyond 1.5 times the interquartile range above the 75th or below the 25th percentile and were excluded prior to analysis. For BNP, donors with NT-proBNP records were excluded and only BNP values were retained to ensure assay consistency.

### Software and computational environment

Metabolic flux inference was performed in R (v4.4.3) using METAFlux (v1.0) with the Human1 model and the OSQP quadratic programming solver (v0.6.0.5)^51^. Pseudobulk aggregation used decoupler (v1.6.0) and scanpy (v1.9.3). Pathway-level statistical analyses (Welch’s *t*-test and Benjamini-Hochberg FDR correction) were performed using scipy (v1.8.0)^55^ and statsmodels (v0.13.2)^56^. Transcription factor regulatory network inference used pySCENIC (v0.12.1)^23^ with arboreto (v0.1.6) for GRNBoost2 and cisTarget motif databases (v10, Aerts Lab). Machine learning classification and regression used scikit-learn (v1.0.2)^57^ and LightGBM (v4.1.0)^58^. Visualization was performed using matplotlib (v3.5.1)^59^ and seaborn (v0.11.2)^60^. All analyses were performed on a macOS system with an Apple M2 Ultra processor (24 cores, 192 GB RAM).

### Code and data availability

Code will be deposited on GitHub upon publication. The Reichart cohort snRNA-seq data are publicly available through CELLxGENE (83ebf054-85ca-48ea-b854- a99756201745). The Chaffin cohort data were obtained from the Single Cell Portal (SCP1303). The Koenig cohort data are available from the Gene Expression Omnibus (GSE183852).

## Acknowledgements

This work was supported by KAKENHI grants from the Japan Society for the Promotion of Science (JSPS) to H.S. (JP23K28184 and JP25H01571) and to S.O. (JP24K03030), JST FOREST Program to H.S. (JPMJFR242Q). Figures were created in part with BioRender.com. We thank T. Suzuoka, Y. Otani, and T. Fujiwara for critical reading of the manuscript, all laboratory members for helpful discussions, and K. Tanaka for assistance with manuscript preparation.

## Author information

**Tomoya Sakuma:** Conceptualization, Methodology, Software, Validation, Formal analysis, Investigation, Data curation, Writing - original draft, Visualization.

**Satoshi Ohno:** Conceptualization, Writing - review & editing, Project administration, Funding acquisition.

**Hideyuki Shimizu:** Conceptualization, Writing - review & editing, Supervision, Funding acquisition.

All authors have read and approved the final manuscript.

## Notes

### Competing Interest Statement

The authors have declared no competing interest.

