## Supplementary Tables S1, S5, S6, S7, S8, Supplementary Figures S1, S2, S3, S4 for "Cell-type-resolved Metabolic Flux Inference Reveals Stromal Metabolic Reprogramming Across Human Cardiomyopathies"

**Supplementary Table 1 | Overview of snRNA-seq datasets**

| | Reichart | Chaffin | Koenig |
|---|---|---|---|
| Reference | Reichart et al.[9] | Chaffin et al.[10] | Koenig et al.[11] |
| Source | CELLxGENE | Single Cell Portal | GEO: GSE183852 |
| Total donors | 78 | 39 | 37 |
| Total nuclei | 869,449 | ~310,000 (DCM) / ~335,000 (HCM) | ~65,000 |
| Healthy donors | 18 | 16 | 25 |
| DCM | 52 | 8 | 12 |
| ACM | 8 | — | — |
| HCM | — | 15 | — |
| Analyzed Genotypes | LMNA, TTN, RBM20, PVneg | TTN, PVneg | — |
| Analyzed cell types | Cardiomyocyte, Endothelial, Fibroblast, Mural, Myeloid | Cardiomyocyte, Endothelial, Fibroblast, Mural, Myeloid | Cardiomyocyte, Endothelial, Fibroblast, Mural |
| Role | Discovery | Validation | Validation |

**Supplementary Table 2 | Complete pathway-level statistics for all 139 Human1 pathways across five cell types**

$Log_2$ fold change (DCM versus healthy donors), Welch's *t*-test statistic, P-value, and FDR for each cell type × pathway combination (n=18 healthy donors, n=52 DCM). Transport and exchange reactions were excluded owing to their limited biochemical interpretability. Provided as CSV.

**Supplementary Table 3 | Three-cohort concordance across the four cell types**

$Log_2$ fold change (DCM versus healthy donors), FDR, and direction concordance for all cell type × pathway combinations across three independent cohorts, restricted to the four cell types shared by all datasets (cardiomyocyte, endothelial, fibroblast, and mural; 4 × 139 = 556 combinations). Provided as CSV.

**Supplementary Table 4 | TF regulon activity changes in DCM.**

Mean AUCell scores (healthy donors and DCM), $log_2$ fold change, and FDR for 128 regulons × 5 cell types (Mann-Whitney U test, donor-level AUCell). Provided as CSV.

**Supplementary Table 5 | Machine learning classification performance (LOOCV, pathway-level features)**

**Healthy donors versus DCM (n=70: healthy donors n=18, DCM n=52)**

| **Model** | **AUC** | **Accuracy** |
|---|---|---|
| SVM linear | 0.999 | 0.971 |
| Ridge | 0.993 | 0.957 |
| ElasticNet | 0.981 | 0.957 |
| LASSO | 0.980 | 0.929 |
| SVM RBF | 0.966 | 0.914 |
| Random Forest | 0.964 | 0.943 |
| LightGBM | 0.902 | 0.857 |

**PVneg versus non-PVneg DCM (n=52: 8 PVneg, 44 Other DCM)**

| Model | AUC | Accuracy |
|---|---|---|
| Random Forest | 0.724 | 0.827 |
| LASSO | 0.500 | 0.846 |
| LightGBM | 0.406 | 0.673 |

Linear/kernel models were excluded due to LOOCV instability under extreme class imbalance (8/52).

**Supplementary Table 6 | LASSO-selected features for healthy donors versus DCM classification**

695 pathway-level features (5 cell types × 139 pathways, transport and exchange pathways excluded) were used as input. LASSO logistic regression with LOOCV selected 15 non-zero features.

| Cell type | Pathway | Coefficient |
|---|---|---|
| Fibroblast | Glycerolipid metabolism | −0.891 |
| Mural | Eicosanoid metabolism | −0.479 |
| Mural | Fatty acid biosynthesis (odd-chain) | −0.443 |
| Myeloid | Pentose and glucuronate interconversions | −0.388 |
| Endothelial | Drug metabolism | +0.317 |
| Fibroblast | Sphingolipid metabolism | −0.218 |
| Myeloid | Propanoate metabolism | +0.150 |
| Fibroblast | Fatty acid desaturation (odd-chain) | −0.120 |
| Myeloid | Phenylalanine metabolism | −0.114 |
| Myeloid | Biopterin metabolism | +0.109 |
| Fibroblast | Fatty acid biosynthesis (unsaturated) | −0.065 |
| Endothelial | Fatty acid oxidation | +0.063 |

| | | |
|---|---|---|
| Mural | Sphingolipid metabolism | −0.059 |
| Myeloid | Beta oxidation of unsaturated fatty acids (n-9, mitochondrial) | +0.045 |
| Fibroblast | Fatty acid elongation (odd-chain) | −0.024 |

**Supplementary Table 7 | Marker gene expression prediction from metabolic pathway scores (LOOCV, DCM n=52)**

| Gene | Best Model | $R^2$ | r | *p* |
|---|---|---|---|---|
| *PPARGC1A* | LASSO | 0.790 | 0.917 | $1.48 \times 10^{-21}$ |
| *MYH7* | Ridge | 0.682 | 0.827 | $4.04 \times 10^{-14}$ |
| *PLN* | Ridge | 0.628 | 0.804 | $6.90 \times 10^{-13}$ |
| *COL1A1* | LASSO | 0.619 | 0.802 | $9.03 \times 10^{-13}$ |
| *ATP2A2* | Ridge | 0.560 | 0.750 | $1.51 \times 10^{-10}$ |
| *ANKRD1* | LASSO | 0.537 | 0.756 | $9.06 \times 10^{-11}$ |
| *COL3A1* | Ridge | 0.456 | 0.678 | $3.39 \times 10^{-08}$ |
| *MYH6* | Ridge | 0.423 | 0.651 | $1.79 \times 10^{-07}$ |
| *POSTN* | LightGBM | 0.292 | 0.550 | $2.40 \times 10^{-05}$ |
| *NPPB* | Ridge | −0.035 | 0.249 | 0.075 |
| *NPPA* | Ridge | −0.044 | 0.290 | 0.037 |

**Supplementary Table 8 | Clinical index prediction from metabolic pathway scores (LOOCV, DCM only)**

| Metric | N | Model | $R^2$ | r | *p* |
|---|---|---|---|---|---|
| LVEF | 43 | Random Forest | −0.041 | 0.103 | 0.513 |
| LVEF | 43 | LASSO | −0.146 | −0.168 | 0.281 |
| LVEF | 43 | Ridge | −0.171 | 0.161 | 0.302 |

| | | | | | |
|---|---|---|---|---|---|
| LVEF | 43 | LightGBM | −0.322 | −0.076 | 0.629 |
| LVIDd | 44 | Random Forest | 0.067 | 0.283 | 0.063 |
| LVIDd | 44 | Ridge | 0.016 | 0.306 | 0.043 |
| LVIDd | 44 | LightGBM | −0.116 | 0.213 | 0.166 |
| LVIDd | 44 | LASSO | −0.125 | 0.086 | 0.579 |
| eGFR | 37 | LASSO | −0.074 | −0.736 | $2.05 \times 10^{-7}$ |
| eGFR | 37 | Random Forest | −0.141 | −0.050 | 0.768 |
| eGFR | 37 | LightGBM | −0.317 | −0.049 | 0.774 |
| eGFR | 37 | Ridge | −0.400 | −0.031 | 0.855 |
| BNP | 21 | Ridge | −0.071 | 0.204 | 0.374 |
| BNP | 21 | Random Forest | −0.102 | −0.040 | 0.863 |
| BNP | 21 | LASSO | −0.141 | −0.080 | 0.729 |
| BNP | 21 | LightGBM | −0.317 | −0.123 | 0.594 |

### Supplementary figure legends

**Supplementary Figure 1 | ATP maximization versus biomass maximization benchmark comparison**

**(A)** Scatter plot comparing cardiomyocyte pathway $\log_2$ fold changes (DCM versus healthy donors) between ATP maximization and biomass maximization objective functions. Each point represents one metabolic pathway. Dashed line indicates the identity line (y = x). Spearman $\rho = 0.165$, $p = 0.053$, indicating low correlation between the two objectives. **(B)** Cell-type flux allocation under maximum ATP production versus maximum biomass production in healthy donors (n=18). Bar heights indicate the percentage of total absolute flux attributed to each cell type (mean ± s.d. across donors). ATP maximization assigned 28.4% of total flux to cardiomyocytes, reflecting the heart's bioenergetic demands, whereas biomass maximization distributed flux nearly equally across all cell types (17.9–22.8%), flattening cell-type-specific metabolic differences. **(C)** Biomass maximization physiological validation heatmap. The same 15 representative pathways from **Fig. 1C** were evaluated under biomass maximization using Z-scores across five cell types. "x" marks pathways where the highest Z-score cell type did not match the biologically expected cell type. Under biomass maximization, 11/15 pathways matched expectations, with cardiomyocyte energy pathways (fatty acid oxidation, TCA cycle, oxidative phosphorylation) and endothelial blood group biosynthesis misassigned. In contrast, ATP maximization achieved 15/15 match (**Fig. 1C**).

**Supplementary Figure 2 | Cross-cohort validation of DCM metabolic alterations and PVneg-specific signatures**

**(A)** Heatmap of $\log_2$ fold changes (DCM versus healthy donor) for four core energy metabolism pathways (oxidative phosphorylation, glycolysis, glycerolipid metabolism, and pyruvate metabolism) across four cell types common to all three cohorts (cardiomyocyte, endothelial, fibroblast, and mural). Columns represent independent cohorts: Reichart (discovery, n=70), Chaffin (validation 1, n=24), and Koenig (validation 2, n=37). All 16 cell type–pathway combinations showed negative $\log_2$ fold changes across all three cohorts. Asterisks denote FDR significance (*FDR < 0.05, **FDR < 0.01, ***FDR < 0.001). **(B)** Dumbbell chart comparing ATP synthesis flux fold changes (genotype mean versus healthy donor mean) between *TTN* and PVneg groups across five cell types in the Reichart (top) and Chaffin (bottom) cohorts. In both cohorts, PVneg exhibited lower ATP flux than *TTN* across all five cell types, reproducing the pattern of greater metabolic suppression in PVneg hearts. **(C)** Heatmap of $\log_2$ fold changes for 24 PVneg-specific pathways (FDR < 0.05 in PVneg only; FDR > 0.1 in all other genotypes in the Reichart cohort). Columns show Reichart PVneg (discovery), Chaffin PVneg (validation), and Chaffin *TTN* (specificity control). PVneg showed concordant direction in 21/24 pathways (88%), whereas *TTN* showed concordance in 18/24 (75%).

**Supplementary Figure 3 | Regulon activity does not differ among DCM genotypes.**

Heatmap of $\log_2$ fold change (versus healthy donor) for the same 25 TF regulons shown in the bipartite network (**Fig. 5D**) across four DCM genotypes (LMNA, TTN, RBM20, PVneg) and five cell types. Kruskal-Wallis tests yielded no significant differences across DCM genotypes (0/125 tests at FDR < 0.05), indicating that transcriptional dysregulation is a genotype-independent feature of DCM.

**Supplementary Figure 4 | Clinical indices of heart failure are not predicted by metabolic pathway scores.**

Scatter plots of observed versus predicted values for three clinical indices using metabolic pathway scores as predictors (LOOCV, DCM donors). **(A)** LVEF (left ventricular ejection fraction; RF, $R^2 = -0.04$, n=43). **(B)** LVIDd (left ventricular internal diameter in diastole; Ridge, $R^2 = 0.02$, n=44). **(C)** eGFR (estimated glomerular filtration rate; LASSO, $R^2 = -0.07$, n=37). All model–metric combinations yielded $R^2 \leq 0.07$, indicating that metabolic pathway scores do not predict hemodynamic severity. This contrasts with accurate disease-state classification (AUC ≈ 1.0; **Fig. 6A**) and marker gene prediction ($R^2 = 0.79$; **Fig. 6D**), suggesting that metabolic remodeling captures molecular pathology rather than clinical severity.

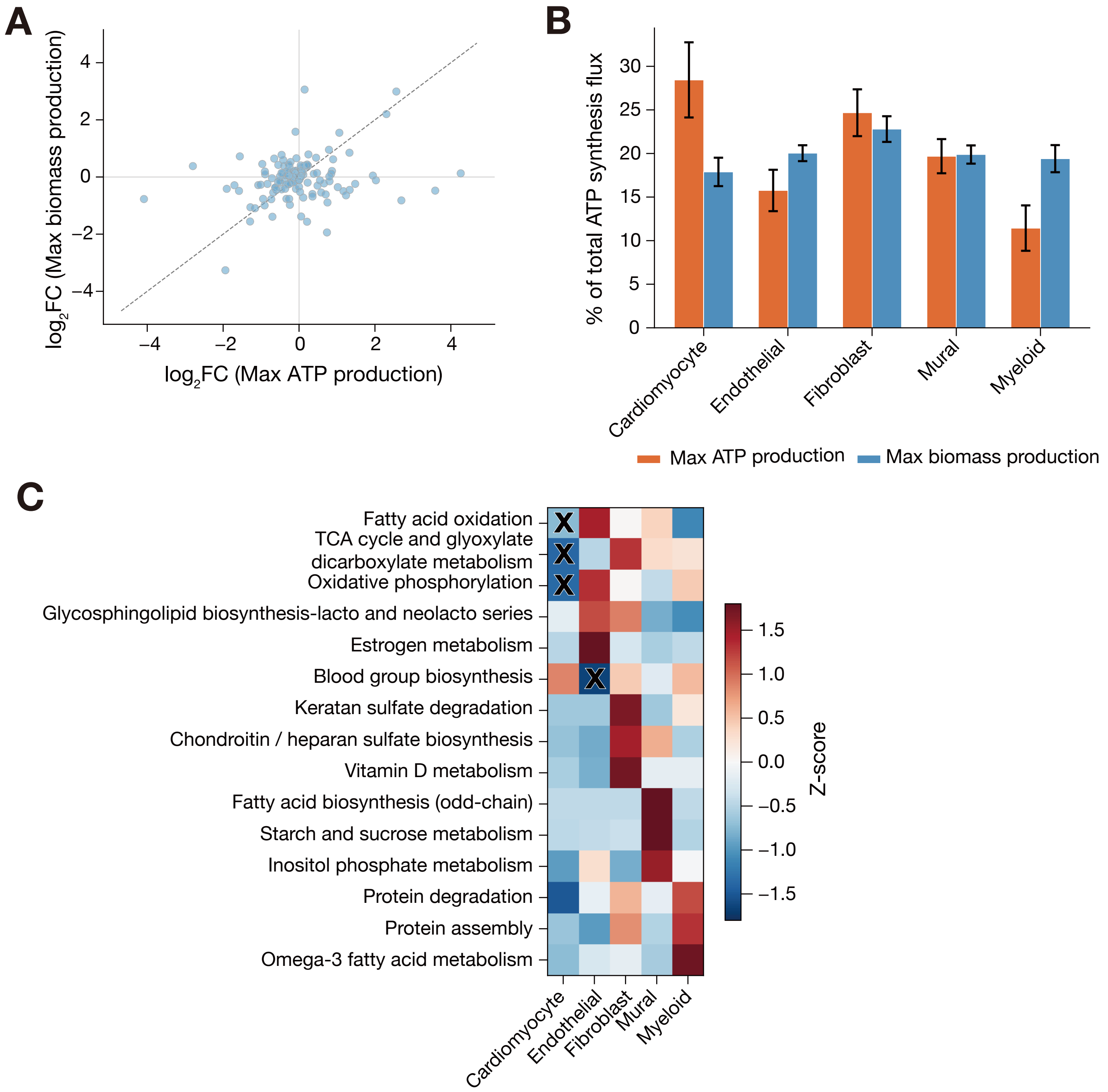


Sakuma et al., Figure S1

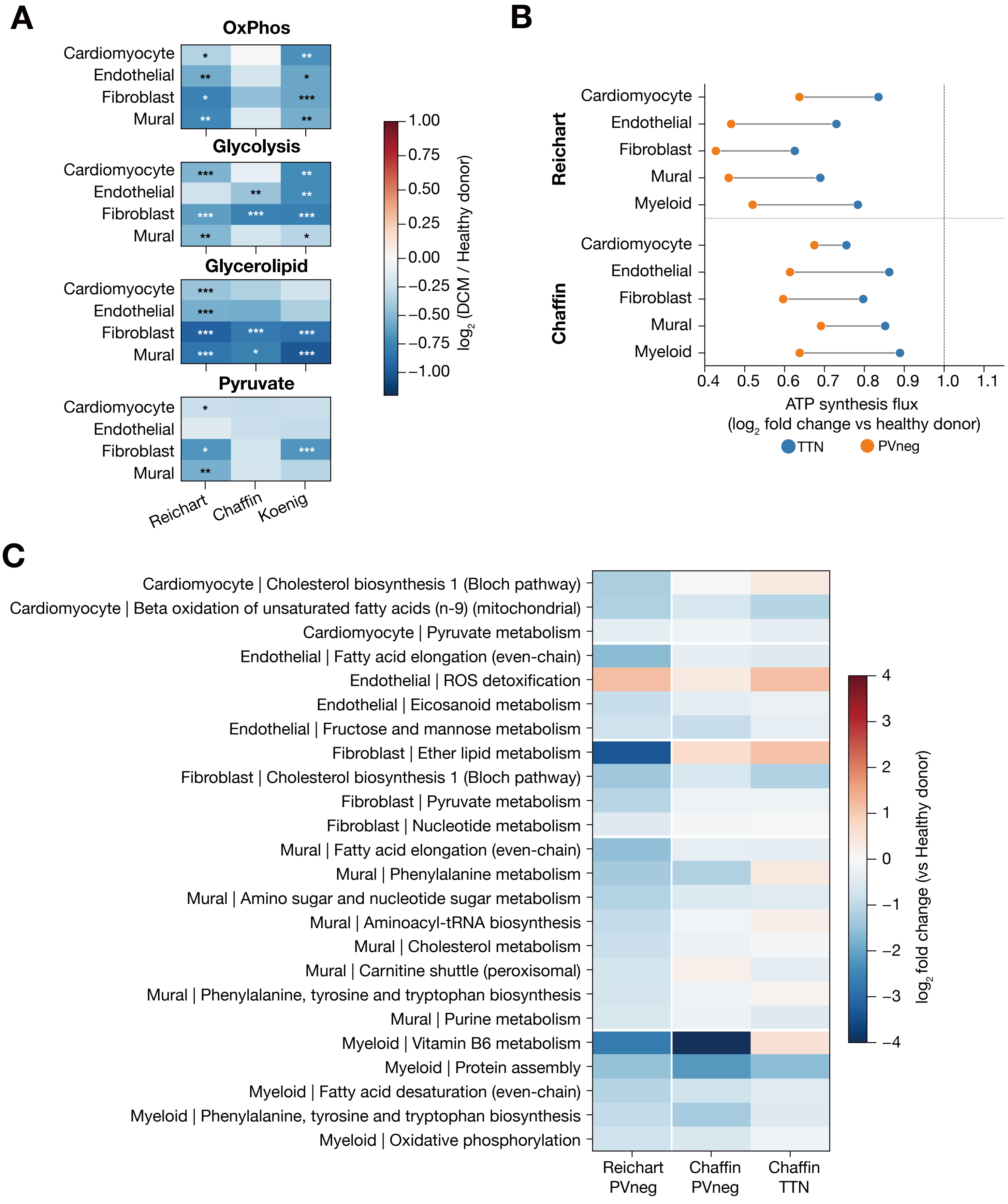


**Sakuma et al., Figure S2**

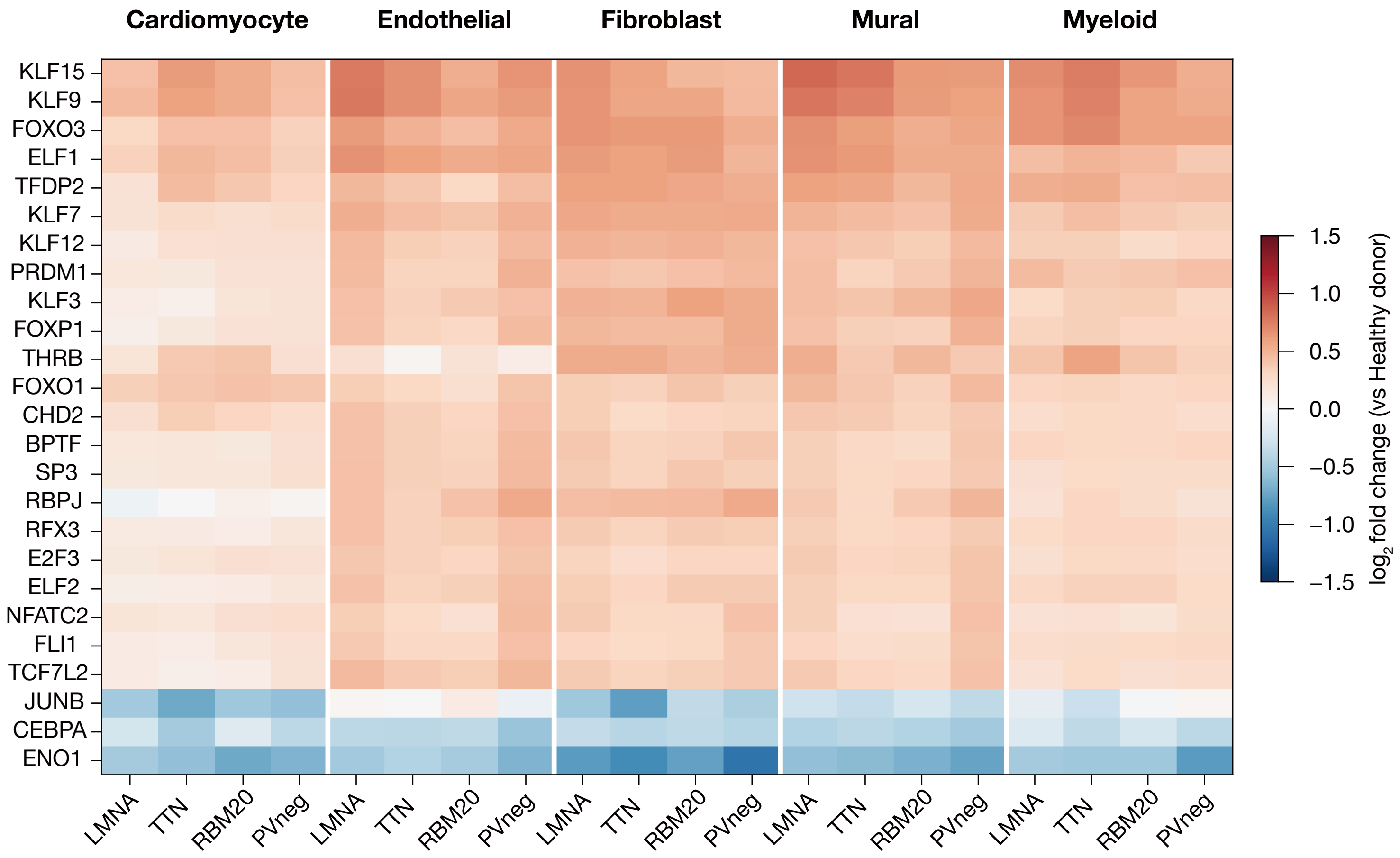


Sakuma et al., Figure S3

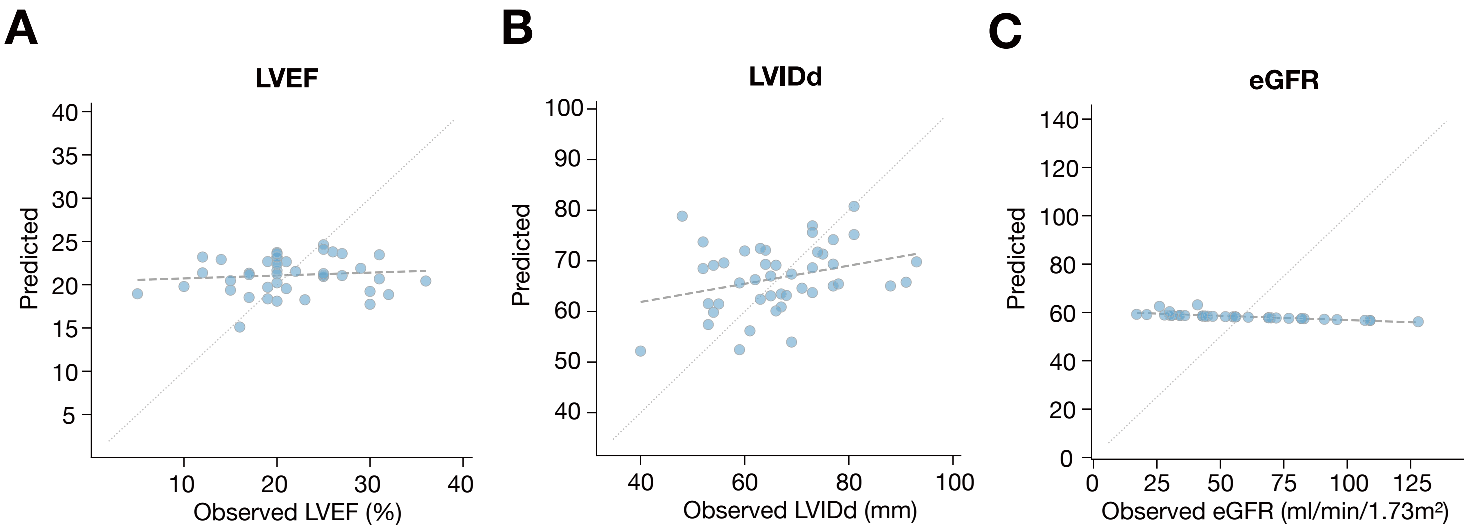


**Sakuma et al., Figure S4**
